# Small molecule modulation of insulin receptor-insulin like growth factor-1 receptor heterodimers in human endothelial cells

**DOI:** 10.1101/2024.03.11.583535

**Authors:** Chloe G Myers, Hema Viswambharan, Natalie J Haywood, Katherine Bridge, Samuel Turvey, Tom Armstrong, Lydia Lunn, Paul J Meakin, Eva M Clavane, David J Beech, Richard M Cubbon, Stephen B Wheatcroft, Martin J McPhillie, Tarik Issad, Colin WG Fishwick, Mark T Kearney, Katie J Simmons

## Abstract

**Objectives:** The insulin receptor (IR) and insulin like growth factor-1 receptor (IGF-1R) are heterodimers consisting of two extracellular α-subunits and two transmembrane β -subunits. Insulin αβ and insulin like growth factor-1 αβ hemi-receptors can heterodimerize to form hybrids composed of one IR αβ and one IGF-1R αβ. The function of hybrids in the endothelium is unclear. We sought insight by developing a small molecule capable of reducing hybrid formation in endothelial cells.

**Methods:** We performed a high-throughput small molecule screening, based on a homology model of hybrid structure. Endothelial cells were studied using western blotting and qPCR to determine the effects of small molecules that reduced hybrid formation.

**Results:** Our studies unveil a first-in-class quinoline-containing heterocyclic small molecule that reduces hybrids by >50% in human umbilical vein endothelial cells (HUVECs) with no effects on insulin or insulin like growth factor-1 receptors. This small molecule reduced expression of the negative regulatory p85α subunit of phosphatidylinositol 3-kinase, increased basal phosphorylation of the downstream target Akt and enhanced insulin/insulin-like growth factor-1 and shear stress-induced serine phosphorylation of Akt. In primary saphenous vein endothelial cells (SVEC) from patients with type 2 diabetes mellitus undergoing coronary artery bypass (CABG) surgery, hybrid receptor expression was greater than in patients without type 2 diabetes mellitus. The small molecule significantly reduced hybrid expression in SVEC from patients with type 2 diabetes mellitus.

**Conclusions:** We identified a small molecule that decreases the formation of IR: IGF-1R hybrid receptors in human endothelial cells, without significant impact on the overall expression of IR or IGF-1R. In HUVECs, reduction of IR: IGF-1R hybrid receptors leads to an increase in insulin-induced serine phosphorylation of the critical downstream signalling kinase, Akt. The underpinning mechanism appears, at least in part to involve the attenuation of the adverse effect of IR: IGF-1R hybrid receptors on PI3-kinase signalling.

**Highlights:**

- We have discovered a small molecule (HI) that inhibits insulin receptor/IGF-1 receptor hybrid formation.
- HI reveals previously unrecognised actions of insulin receptor/IGF-1 receptor hybrids distinct to insulin and IGF-1 receptors in endothelial cells.
- Treatment of endothelial cells with HI enhances activity of the downstream signalling kinase Akt due to inhibitory regulation via PI3-K.

## 1. Introduction

Over the past four decades, changes in human lifestyle have contributed to an explosion of obesity[1, 2] and its frequent sequela, type 2 diabetes mellitus (T2DM)[3]. A poorly understood hallmark of obesity and T2DM is insulin resistance [4, 5], leading to dysregulation of cellular growth and nutrient handling[6]. The insulin receptor (IR) acts as a conduit for insulin-encoded information, which is transferred via a complex intracellular signalling network including the critical signalling nodes phosphatidylinositol 3-kinase (PI3-K) and the serine/threonine kinase Akt, to regulate cell metabolism[7]. During evolution, the IR and insulin-like growth factor-1 receptor (IGF-1R) diverged from a single receptor in invertebrates[8, 9], into a more complex system in mammals[10]. Stimulation of IR or IGF-1R initiates phosphorylation of IR substrate (IRS) proteins at multiple tyrosine residues[7], which can then bind PI3-K, initiating the conversion of the plasma lipid phosphatidylinositol 3,4,-bisphosphate to phosphatidylinositol 3,4,5-trisphosphate (PIP3), which activates the multifunctional serine−threonine kinase Akt [11].

The endothelium plays an important role in cardiovascular health and diseases. In addition to controlling blood homeostasis and fibrinolysis, it regulates vascular tone and promotes angiogenesis. In endothelial cells (EC), Akt activates the endothelial isoform of nitric oxide synthase (eNOS) by phosphorylation of Ser1177[12, 13]. In humans and other mammals, despite high structural homology and activation of similar downstream pathways, the biological processes regulated by insulin and IGF-1 are strikingly different[14]. Consistent with this, in EC, we demonstrated that deletion of IR reduced[15, 16], whereas deletion of IGF-1R increased, basal ^Ser1177^peNOS and insulin-mediated ^Ser1177^peNOS[17]. More recently, we showed that increasing IR in EC enhances insulin-mediated ^Ser473^pAkt but blunts insulin-mediated ^Ser1177^peNOS[18], whereas increased IGF-1R reduces basal ^Ser1177^peNOS and insulin-mediated ^Ser1177^peNOS[19].

IR and IGF-1R are homodimers consisting of two extracellular α-subunits and two transmembrane spanning β-subunits[20]. IR and IGF-1R can heterodimerize to form ‘hybrid’ receptors composed of one IR αβ complex and one IGF-1R αβ complex[21]. In skeletal muscle, fat and the heart, hybrid expression has been shown to exceed that of IGF-1R and IR[22]. While the role of hybrids in human physiology is undefined, there is a clear association with increased hybrids and pathological conditions of metabolic stress including: T2DM[23, 24], obesity[25], hyperinsulinemia[26], insulin resistance[27] and hyperglycaemia[28]. Hybrid receptors are not amenable to genetic manipulation; therefore, development of small molecules to inhibit hybrid formation is an attractive approach to examine their physiological role. Here, we describe the rational, structure-based discovery of a small molecule that inhibits formation of hybrids in human cells.

## 2. Methods

### 2.1 Hybrid Homology Model Generation

The homology model of the hybrid ectodomain was based on the IR ectodomain dimer[29]. The IGF-1R ‘monomer’ was created using I-TASSER[30]. The IGF-1R monomer was merged with the IR monomer from the crystal structure and the resulting dimer prepared using the Maestro graphical user interface (Schrodinger, LLC, New York, NY, 2017), using the protein preparation wizard to minimise steric clashes between residues and create disulfide links between the two monomers (**Figure S1**).

### 2.2 Virtual-High Throughput Screening

The ZINC Database of drug-like molecules was screened on an ARC3 high-performance computing cluster. This library was docked to the hybrid homology model using Glide high-throughput virtual screening mode[31]. A subset of the 100,000 top scoring molecules was re-scored using Glide Standard precision mode. The top 5000 scoring molecules were re-docked using an alternative docking software, AutoDock Vina[32]. The 5000 molecules were clustered by energy (predicted binding affinity) and an arbitrary cut-off of −7.5 chosen for visual inspection. Compounds that satisfied these criteria were cross-referenced with the top-ranking molecules from Glide to identify those which were predicted to bind favourably in both docking softwares. Additional *in silico* analysis was performed using Maestro and the in-house *de novo* design software SPROUT[33] to identify compounds predicted to make hydrogen bonds and Van der Waals interactions with the IR and steric clashes with the IGF-1R.

### 2.3 Ligand-Based Virtual Screening

A shape similarity screen was performed using OpenEye’s ROCS software[34, 35]. ROCS allows rapid identification of active compounds by shape comparison, through aligning and scoring a small molecule database against a query structure. The EON software compares electrostatic potential maps of pre-aligned molecules to a query structure, producing a Tanimoto similarity score for comparison. The initial hit molecule query file was the docked pose from the initial virtual screening run and the library used was the University of Leeds’ Medicinal Chemistry and Chemical Biology library. The library contains around 27,000 molecules. Standard parameters were used, as well as an EON input command. The top 1000 ranked molecules were taken into EON (electrostatic matching) and the top 100 molecules visually inspected using the VIDA graphical user interface. Molecules were selected for biological evaluation based on favourable shape and electrostatic matching to HI-2.

### 2.4 Human Cell Line Culture

Human embryonic kidney (HEK293) cells were cultured in Dulbeccoʹs Modified Eagleʹs Medium (DMEM, Merck) containing 10% Fetal Calf Serum and 1% Antibiotic Antimycotic Solution (Sigma Aldrich). Human umbilical vein endothelial cells (HUVECs, PromoCell) were cultured in Endothelial Growth Medium-2 (ECGM-2, PromoCell) containing relevant growth supplements (PromoCell) and 1% Antibiotic Antimycotic Solution. Cells were maintained at 37°C, 5% CO_2_.

### 2.5 Bioluminescence Resonance Energy Transfer (BRET) Assay

HEK293 cells were seeded at a density of 200,000 cells per well in a six-well plate and transfected with IR-A-Rluc only (control) or IR-A-Rluc and IGF-1R-YPET (hybrids) using Lipofectamine 2000, as the transfection reagent and incubated at 37°C and 5% CO_2_ [36]. This results in the biosynthesis of three populations of receptors: (IR-A-Rluc)_2_ homodimers, (IGF-1R-YFP)_2_ homodimers, and IR-A-Rluc/IGF-1R-YFP heterodimers (**Figure S2**). Since only hybrids can produce a BRET signal, this method allows direct study of hybrids without having to separate them from homodimers of each type. One day after transfection, cells were transferred into 96-well microplates (white, Poly-D-Lysine coated, Perkin Elmer) at a density of 30,000 cells per well. Forty-eight hours after transfection, cell culture medium was removed and replaced with media containing compound of interest (100 µM) or vehicle control (dimethyl sulfoxide (DMSO, equivalent to 100 µM concentration). After a further 24 hours, cells were washed with 50 μl of phosphate buffered saline (PBS, Sigma Aldrich) prior to Bioluminescence Resonance Energy Transfer (BRET) measurements. BRET measurements were taken in PBS (final volume 50 μl) containing coelenterazine (stock solution (1 mM) in ethanol, final concentration 5 µM). BRET measurements quantify Rluc light emission (485 nm) and YPET light emission (535 nm) using a PerkinElmer microplate reader. BRET signal was calculated using the following equation:

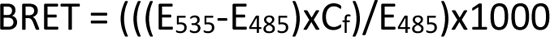

where C_f_= E_535_/E_485_ for the IR-Rluc only transfected HEK293 cells.

To validate our assay, we showed increased BRET signal when cells were treated with IGF-1 (100 nM) compared to basal levels, as described by Blanquart et. al.[37] (**Figure S3**). Based on these assays the HI series was taken forward for further in-vitro experiments.

### 2.6 LIVE/DEAD Cell Viability Assay

Quantification of cell viability was performed using the LIVE/DEAD Assay Kit (Thermo Fisher Scientific), as per manufacturer’s protocol with minor modifications. Briefly, HEK293 cells were seeded at a density of 60,000 cell per well, in a 96-well poly-L-lysine-coated plate (Corning) and incubated overnight at 37 °C with 5% CO_2_. After incubation, the desired compound was added to fresh media (DMEM, Merck) at a concentration of 100 µM before being added to the cells and incubated for 24 hours. After 24 hours, media containing the compound was removed and cells washed in PBS (200 µl). Twenty microliters of 2 mM EthD-1 stock solution was added to 10 mL of PBS to give a 4 µM EthD-1 solution. Five microliters of 4 mM calcein AM stock solution was then added to the 10 mL EthD-1 solution. One hundred microliters of the combined LIVE/DEAD® assay reagent was added to each well and incubated for 45 min before quantification of percentage of red and green cells using the IncuCyte® ZOOM (Sartorius) system and its proprietary ZOOM 2016A software.

### 2.7 HI-2 Inhibitor Treatment of Endothelial Cells

HUVECs were seeded at a density of 200,000 cells per well, in a six-well culture plate and maintained at 37°C and 5% CO_2_ until 90-95% confluent, before they were treated with HI-2. Cells were washed with pre-warmed PBS and 1ml of fresh ECGM-2 media (PromoCell) containing HI-2 at final concentration of 100 µM (or vehicle control DMSO, Sigma-Aldrich) was added to the cells. Cells were incubated 24 hours prior to cell lysis.

### 2.8 HI-2 Pre-treatment of Endothelial Cells with Insulin or IGF-1

HUVECs at passage 3 to 6 were cultured in a six-well plate in ECGM-2, as described above. Ninety percent confluent HUVECs were washed with pre-warmed PBS and replaced with 0.2% FCS-containing basal media, overnight before stimulation with insulin (25, 50, 100 nM, Sigma Aldrich) or IGF-1 (25, 50, 100 nM, GroPep Bioreagents), at different doses for 10 minutes, lysed and harvested for Western blot analysis. Prior to lysis, cells were washed twice in 2mL ice-cold PBS. A 100μL aliquot of Cell Extraction Buffer (Invitrogen) containing 1μL/mL protease inhibitor solution (Invitrogen) was added to each well and left on ice for 5 minutes. Cells were scraped and collected and left on ice for 10 minutes, with intermittent vortexing. A 20-minute centrifugation step at 4°C and 13,000g was performed, collecting the supernatant into fresh receptacles. Protein quantification was performed using the Pierce™ BCA Protein Assay kit (Thermo Fisher Scientific), as per manufacturer’s protocol.

### 2.9 Immunoprecipitation

A solution containing 300μL of binding buffer (100 mM NaCl; 10 mM MgSO_4_; 100 mM HEPES; 0.025% Tween-20; pH 7.8), 1μL/mL protease and phosphatase inhibitors, and 30μL Protein G Agarose beads (Sigma Aldrich) was combined with 3μL of primary antibody and 150 μg of total cell lysate. Samples were incubated at 4°C for 16-18 hours, medium speed on a MACSmix™ Tube Rotator (Miltenyi Biotec). Upon completion, samples were centrifuged for one minute at 500g, collecting the supernatant into a separate fresh receptacle. The pellet was washed six times in 1mL of PBS-T (0.05% Tween-20) containing 1μL/mL protease and phosphatase inhibitors. Thirty microliters of combined buffers of 1X SDS NuPAGE™ LDS and 1X Reducing Agent (Invitrogen) were added to the beads. Prior to loading, samples were incubated at 95°C for five minutes and centrifuged for one minute at 1000g to pellet the beads. The supernatant was then used for SDS-PAGE.

### 2.10 Immunoblotting

Immunoblotting was carried out using previously published methods[39]. Target proteins were probed using their respective antibodies (**Table S1**) in 5% Bovine Serum Albumin (BSA, Sigma-Aldrich) overnight at 4°C. The membrane was further incubated with horseradish peroxidase (HRP)-conjugated anti-mouse or anti-rabbit secondary antibodies (1:5,000) in 5% milk buffer for 1 hour at room temperature. Blots were visualised using Immobilon® Western Chemiluminescent HRP Substrate reagents (Millipore), using the Gel documentation system, Syngene G:BOX.

### 2.11 Real-Time Polymerase Chain Reaction

Real-time PCR was carried out using previously published methods[39] using the following primers: IGF1R - PrimePCR™ SYBR® Green Assay: IGF1R, Human (BioRad-assay ID qHsaCED0044963) INSR - PrimePCR™ SYBR® Green Assay: INSR, Human (BioRad-assay ID qHsaCID0017132) and ACTB - PrimePCR™ SYBR® Green Assay: ACTB, Human (BioRad-assay ID qHsaCED0036269). The cycles to threshold was measured for each well, the average of triplicate readings for each sample taken, normalised to ACTB.

### 2.12 Akt Activity Assay

Akt kinase activity was analyzed by nonradioactive immunoprecipitation-kinase assay, as previously described[38], according to manufacturer’s protocol (Cell Signaling Technology; Boston, MA). Cell extracts of 30 µg were incubated with immobilized Akt 1G1 monoclonal antibody. After extensive washing, the kinase reaction was performed at 30°C for 30 minutes in the presence of 200 µM cold ATP and GSK-3 substrate. Phospho-GSK-3 was measured by Western blot, using phospho-GSK-3α/β (Ser-21/9) antibody, provided in the assay kit.

### 2.13 Endothelial Cells exposure to Shear Stress

Using previously published methods[39], HUVECs were seeded at a density of 200,000 cells per well, into 6-well plates and incubated at 37°C, 5% CO_2_. Once 75% confluent, monolayers were placed onto an orbital rotating platform (Grant Instruments) housed inside the incubator. The radius of orbit of the orbital shaker was 10 mm, and the rotation rate was set to 210 rpm for 24 hours, which generated a swirling motion of the culture medium over the cell surface volume. The movement of fluid due to orbital motion represents a free surface flow at the liquid–air interface at 12 dyne/cm. Cells were harvested in 100 µL of Cell Extraction Buffer containing 1 μL/mL protease inhibitor solution (Invitrogen) for protein analysis.

### 2.14 siRNA-mediated knockdown of IR and IGF-1R

Reduction of hybrid receptor expression in HUVECs was carried out by transfection of validated human siRNA duplexes of insulin receptor (Silencer™ Pre-Designed siRNA, Invitrogen; Catalog number: AM51331, siRNA ID:29) and IGF-1 receptor (Catalog number: AM51331, siRNA ID:110754) with Lipofectamine RNAiMax Transfection Reagent (Invitrogen), according to manufacturer’s established protocol, as we previously reported[39]. Scrambled siRNA (Invitrogen; Catalog number: AM4613) was used as negative control (Invitrogen) for transfection. Briefly, HUVECs were rendered 70% confluent and transfected with 25 pmol siRNA for 24 hours, before cell lysates were prepared for western blotting analysis.

### 2.15 Isolation of saphenous vein endothelial cells

SVECs were obtained from consenting patients undergoing coronary artery bypass graft surgery, at the Leeds General Infirmary (Ethical Approval CA01/040) and were isolated using previously described methods[40]. Confluent SVEC at passage 3 to 7 were used for experiments and analysis.

### 2.16 High-Performance Liquid Chromatography Solubility Assay

This method was adapted from Bellenie et. al[41]. Briefly, 10 µL of 10 mM DMSO stock solution was pipetted into a micro-centrifuge tube (Sarstedt) containing 990 µL of PBS buffer (pH 7.4, Sigma Aldrich) and mixed for 5 seconds on vortex mixer to create 100 µM solution of compound with 1% DMSO. Following agitation of the suspension at 500 rpm for 2 hours at 20°C, it was separated by centrifugation (14000 rpm, 15 min). Two hundred microliters of the supernatant was transferred to a vial containing 50 µL of DMSO (Sigma-Aldrich) and mixed for 5 seconds to avoid precipitation from the saturated solution. The concentration of the solubilized compound in solubility sample is measured by high performance liquid chromatography (HPLC) with UV detection, using an external standard, prepared by pipetting 10 µL of the same batch of compound-DMSO stock used in solubility sample preparation into 990 µL of DMSO. The detail of the HPLC method is as follows: chromatographic separation at 30°C was carried out over a 5 minute gradient elution method from 90:10 to 10:90 water: methanol (both modified with 0.1% formic acid) at a flow rate of 1.5 mL/min. A calibration curve was prepared by injecting 0.5, 2.5, and 5 µL of compound external standard. Compound solubility value was obtained by injecting 2 and 20 µL of sample.

### 2.17 Statistics

Results are expressed as mean±SEM. Comparisons were made using paired Student t tests or unpaired Student t tests, as appropriate using Prism Software version 9 (GraphPad). P<0.05 was considered statistically significant. Significant P values are shown as:*P<0.05,**P<0.01,***P<0.001, and****P<0.0001

## 3. Results

### 3.1 Virtual screening to identify potential small-molecule inhibitors of insulin receptor-insulin like growth factor receptor hybrids

We generated a homology model of the hybrid receptor ectodomain (**Figure S1A**) based on the IR ectodomain dimer described by Croll *et al*[29]. We examined this hybrid homology model using the KFC2 protein-protein interaction hotspot prediction algorithm[42] to identify key regions involved in the IR/IGF-1R protein-protein interaction (**Figure S1B**). Each of three hotspots identified was evaluated to determine their suitability for docking studies. We selected a hotspot covering amino acids 400-570 in IR for virtual high-throughput screening (**Figure S1C)**. This region had the lowest degree of sequence identity between IR and IGF-1R, affording us the greatest likelihood of identifying a selective modulator of hybrid formation. Using a library of commercially available small molecules and Glide docking software [31] we identified a set of 5000 ligands binding to insulin receptors in a way they could prevent hybrid formation. The best scoring molecules were further examined using our in house *de novo* design program SPROUT[33] **(Figure S1D)**.

We purchased 42 compounds identified from our library of seven million compounds (**Table S2**) from commercially-available libraries and used liquid chromatography-mass spectrometry (LCMS) to determine compound purity and integrity. A BRET assay, measuring the ratio of IR-Rluc interacting with IGF-1R-YPET-a means of quantifying hybrid levels specifically, was used to find a potential candidate inhibitor of hybrid formation. Amongst these, compound HI-1 (**Figure 1A**) reduced hybrid formation in the BRET assay by ∼20 % (**Figure 1B**). HI-1 is predicted to bind to IR and make key hydrogen bonding interactions with Gly463, Ala466 and Arg577, but crucially is predicted to make a steric clash with Arg450 and Asn451 of IGF-1R-specifically preventing hybrid formation (**Figure 1C**). A LIVE-DEAD Cell Viability Assay was used to examine compound toxicity in HEK293 cells (**Figure 1D**). Compound HI-1 had some toxicity compared to cells treated with vehicle control (DMSO). To circumvent this, we developed a series of analogues of HI-1 from both in-house synthesis (**43-56, Table S3**) and commercial sources (**57-63, Table S3**). Compounds were synthesised using the schemes described in **Schemes S1-S3**. Briefly, 3-nitro-1*H*-1,2,4-triazole is coupled with aryl alkyl chlorides before reduction of the nitro-group to give the free amine. A small molecule X-ray crystal structure of the nitro-triazole intermediate (**Figure S4, Tables S5-10**) confirmed the regioselectivity of the N-alkylation reaction. Ring-expansion of substituted isatins using the Pfitzinger reaction gave the 4-quinolinecarboxylic acid derivatives, which are then coupled to the amino-triazole derivatives using propylphosphonic anhydride.

**Figure 1.**
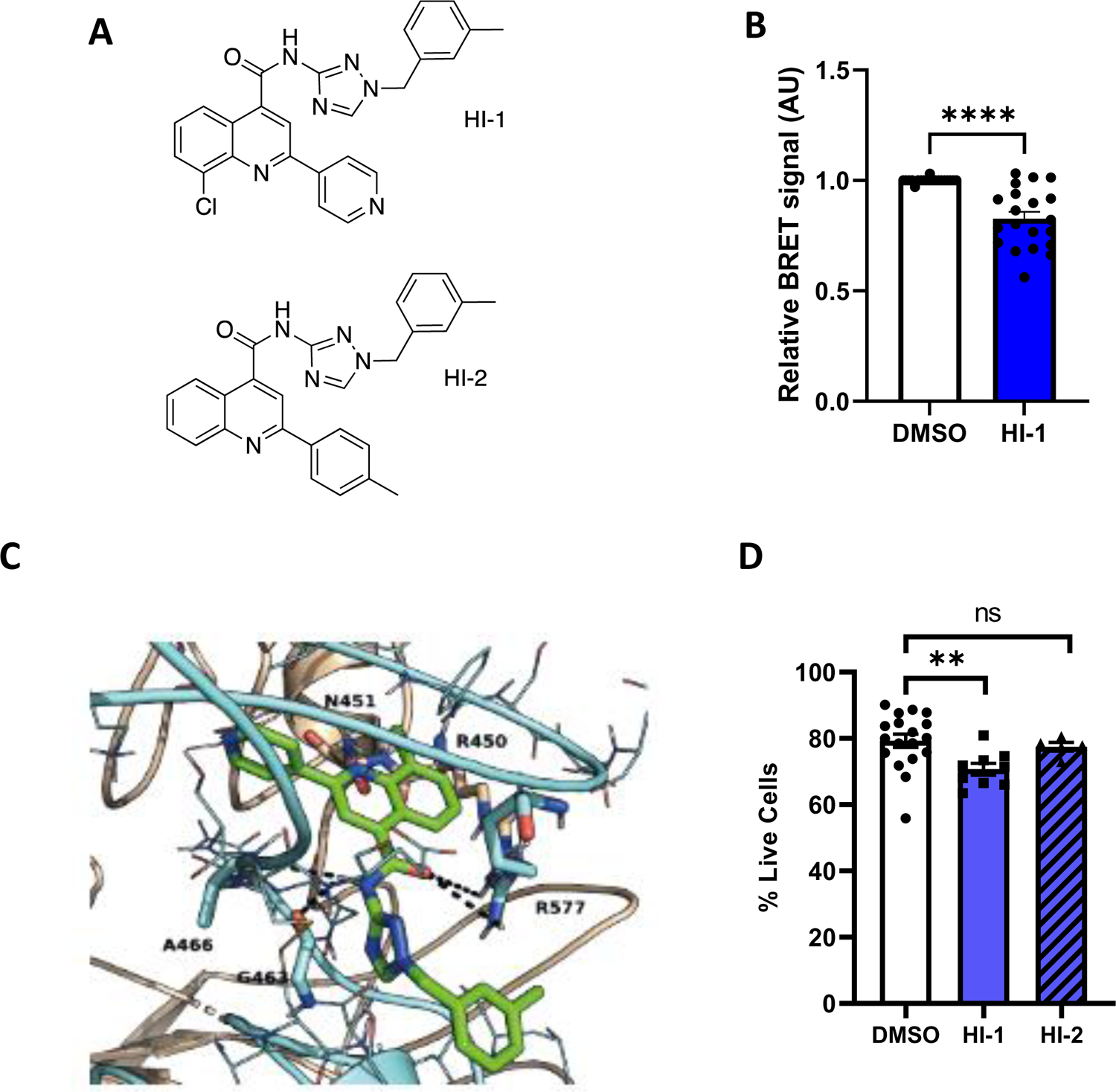
Identification of potential small-molecule inhibitors of insulin receptor-insulin like growth factor receptor hybrids. A) Structures of compounds hybrid inhibitor 1 (HI-1) and hybrid inhibitor 2 (HI-2). B) Compound HI-1 from the virtual high-throughput screen significantly reduced hybrid formation in BRET assay performed in HEK293 cells compared to vehicle control (250 µM, 24h treatment, n=8,3; p= <0.0001). C) Predicted binding mode of HI-1 to IR (wheat ribbons), making key hydrogen bonding interactions with Gly463, Ala466 and Arg577, but crucially makes a steric clash with Arg450 and Asn451 of IGF1R (cyan ribbons)-specifically preventing hybrid formation. A number of residues involved in the interaction are highlighted as wheat/cyan sticks. Image generated using Pymol. D) Analysis of compound toxicity in HEK293 cells for the HI series showed that HI-1 significantly reduced cell viability, whereas there was no effect of HI-2 compared to vehicle control (DMSO) (100 µM, 24h treatment, n=5, p=0.0103 for HI-1 and p=0.2315 for HI-2). All values are expressed as mean and standard error of the mean (SEM). Statistical significance was determined by the Student’s t-test. ns denotes non-significant differences found.

These compounds were screened using our BRET assay (**Figure S5**) and the most active of these compounds, is henceforth described as HI-2 (**Figure 1A**). Due to its structural similarity to our HI series, we also tested the neurokinin 3 receptor antagonist, talnetant (**64**). However, this molecule showed no effect on hybrid formation (**Figure S3**). We used the ligand-based screening tool ROCS[34, 35] and identified compounds from our in-house library of 30,000 ligands which matched the shape and electrostatic properties of the HI series, but with a different central pharmacophore (**65** & **66, Table S3**). Screening results for these compounds showed that it was crucial to retain the quinoline-core for activity (**Figure S5**).

### 3.2 Effect of HI-2 on expression of hybrids in HUVECs

In HUVECs expressing both isoforms of the insulin receptor IR-A and IR-B[44], HI-2 showed no significant cell viability effects compared to vehicle control (**Figure 1D**) and was more potent than HI-1, reducing hybrid expression by ∼50% in HUVECs (**Figure 2A**). This was quantified using the technique first described by Li *et al*.[43] applying HI-2 at 100µM concentration. To estimate the relative abundances of homo-IR, hybrids, and homo-IGF-1R, cell lysates were immunoprecipitated with anti-IGF-1R antibody and the levels of IR and IGF-1R in immunoprecipitate, supernatant and cell lysate not subjected to immunoprecipitation was performed. The IR in the immunoprecipitate represented IR from hybrids, whereas IR remaining in supernatant were likely homo-IR/IR. HI-2 reduced hybrid expression, with no effect on protein expression of IR (**Figure 2B**) or IGF-1R (**Figure 2C**) protein levels. Expression of the genes encoding IR and IGF-1R were also unchanged in HI-2 treated HUVECs (**Figure 2D & E**).

**Figure 2.**
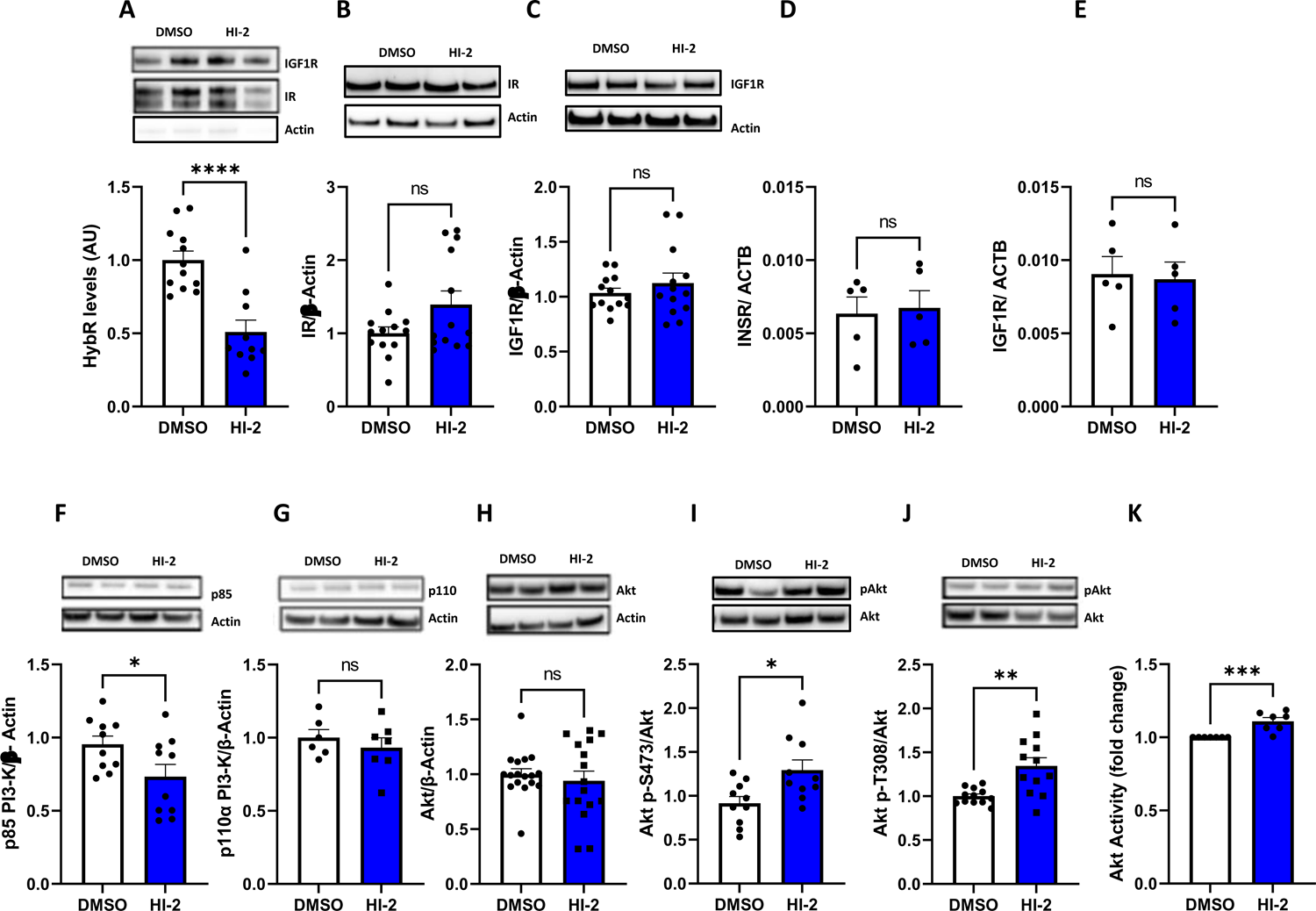
Hybrid receptor inhibition by HI-2 mediates activation of Akt. A) Expression of hybrid formation in human umbilical vein endothelial cells (HUVECs) treated with HI-2 for 24h compared to vehicle control (DMSO). To estimate the relative levels of hybrids, cell lysates were immunoprecipitated with anti-IGF1R antibody, and the levels of IR and IGF1R in immunoprecipitation, supernatant, and cell lysate before immunoprecipitation determined. The IR in immunoprecipitation represented IR from hybrids, whereas IR remained in supernatant were likely homo-IR/IR. (100 µM, 24h treatment, n=10 p= <0.0001). B) IR dimer formation in HUVECs treated with HI-2 compared to vehicle control (100 µM, 24h treatment, n=13 p= 0.07). C) IGF1R dimer formation in HUVECs treated with HI-2 compared to vehicle control (100 µM, 24h treatment, n=13 p= 0.3867). D) Expression of the gene encoding IR in HUVECs treated with HI-2 compared to vehicle control (100 µM, 24h treatment, n=5 p= 0.8141). E) Expression of the gene encoding IGF1R in HUVECs treated with HI-2 compared to vehicle control (100 µM, 24h treatment, n=5 p= 0.8396). F) Total p85 PI3-K protein expression in HUVECs treated with HI-2 compared to vehicle control (100 µM, 24h treatment, n=10 p= 0.0417). G) Total p110α PI3-K protein expression in HUVECs treated with HI-2 compared to vehicle control (100 µM, 24h treatment, n=5 p= 0.1879). H) Akt protein expression in HUVECs treated with HI-2 compared to vehicle control (100 µM, 24h treatment, n=16 p= 0.5581). I) Ser473-pAkt protein in HUVECs treated with HI-2 compared to vehicle control (100 µM, 24h treatment, n=10 p= 0.0142). J) Thr308-pAkt in HUVECs treated with HI-2 compared to vehicle control (100 µM, 24h treatment, n=12 p= 0.0019). K) Akt activity in HUVECs treated with HI-2 compared to vehicle control (100 µM, 24h treatment, n=7 p=0.0009). All values are expressed as mean and standard error of the mean (SEM). Statistical significance was determined by the Student’s t-test. ns denotes non-significant differences found. DMSO is vehicle control.

### 3.3 Effect of HI-2 on basal expression of key insulin signalling nodes in HUVECs

We then examined the effect of HI-2 on basal expression of molecules at the critical signalling nodes in the insulin/IGF-1 signalling pathway; PI3-K and the serine/threonine protein kinase Akt. After treatment with HI-2 there was a >20% reduction in expression of the p85α negative regulatory subunit of PI3-K (**Figure 2F**) but no change in the catalytic subunit p110α (**Figure 2G**). HI-2 had no effect on basal expression of Akt (**Figure 2H**). HI-2 led to a ∼40% increase in ^Ser473^pAkt (**Figure 2I**) and ^Thr308^pAkt (**Figure 2J**) and an increase in Akt activity (**Figure 2K**).

### 3.4 Impact of HI-2 on endothelial cell insulin and IGF-1 signalling

We investigated the effect of HI-2 on insulin-mediated phosphorylation of Akt. Insulin-induced ^Ser473^pAkt was increased approximately three-fold across a range of insulin concentrations after treatment with HI-2 for 24 hours (**Figure 3A**). The increase in ^Ser473^pAkt was blunted when cells were treated with the PI3-K inhibitor, Wortmannin (**Figure 3B**); suggesting that this effect is mediated via the PI3-K-dependent pathway. Additionally, IGF-1 induced ^Ser473^pAkt was increased to a similar level as insulin across a range of IGF-1 concentrations after treatment with HI-2 for 24 hours (**Figure 3C**).

**Figure 3.**
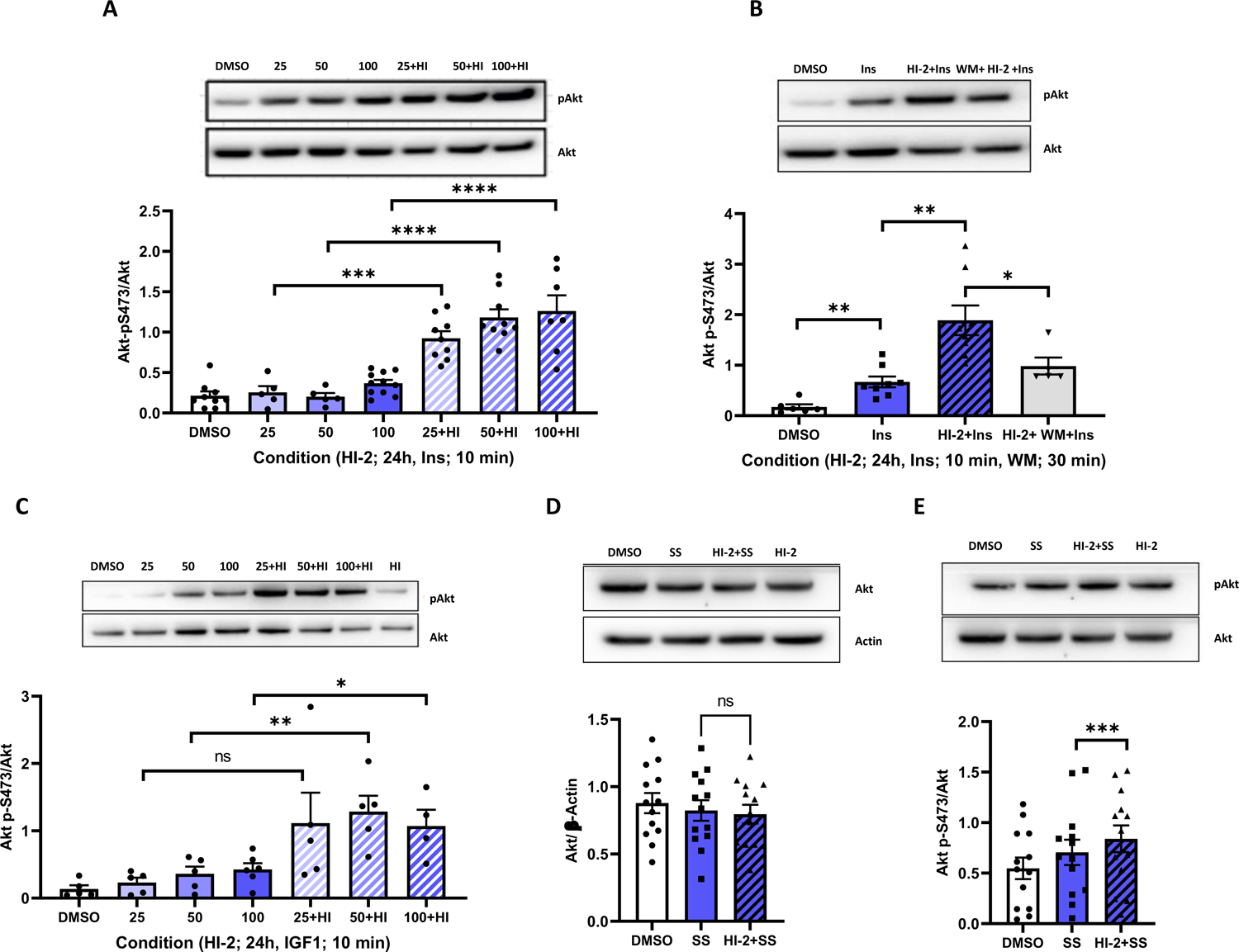
Effects of HI-2 on endothelial cell insulin and IGF-1 signalling. A) Insulin-induced Ser473-pAkt in HUVECs treated with HI-2 compared to vehicle control (100 µM HI-2 for 24h at varying insulin concentrations (Ins, 25, 50, 100nM) for 10 minutes, n=>5, p= 0.003 (25 nM), <0.0001 (50 nM), 0.0033 (100 nM). B) Insulin-induced Ser473 pAkt in HUVECs treated with HI-2 and the PI3-K inhibitor Wortmannin (100 µM HI-2 for 24hours, 100nM insulin and wortmannin, WM, 50 nM; 30 minutes, n=>5, p= 0.0027 (DMSO vs Insulin), 0.0016 (Insulin vs Insulin + HI-2), 0.0443 (Insulin + HI-2 vs Insulin + HI-2 + Wortmanin). C) IGF1-induced Ser473 pAkt in HUVECs treated with HI-2 compared to vehicle control (100 µM HI-2 for 24 h at varying IGF1 concentration (IGF1 25, 50, 100nM) for 10 minutes, n=>4, p= 0.0910 (25 nM), 0.007 (50 nM), 0.0204 (100 nM). D) Shear stress-induced (SS) changes in total Akt protein in HUVECs treated with HI-2 (100 µM, 24 h treatment, shear stress for 24 h, n=13 p= 0.7987). E) Shear-induced Ser473 pAkt in HUVECs treated with HI-2 compared to vehicle control (100 µM, 24h treatment, shear stress for 24 h, n=13, p=0.4693). All values are expressed as mean and standard error of the mean (SEM). Statistical significance was determined by the Student’s t-test. ns denotes non-significant differences found. DMSO is vehicle control.

### 3.5 Effect of HI-2 on shear stress mediated phosphorylation of Akt in endothelial cells

As described previously[12, 13], Akt is activated by shear stress, independent of ligand binding to cell surface receptors. This has been shown to be PI3-K-dependent. We therefore, examined the possibility that HI-2 having an impact on shear-induced phosphorylation of Akt. In these experiments HI-2 had no effect on the expression of total Akt (**Figure 3D**). However, HI-2 increased shear-induced ^Ser473^pAkt by ∼20% (**Figure 3E**).

### 3.6 Reducing hybrids blunts the effect of HI-2 on Akt phosphorylation

To explore the effect of reducing hybrids in endothelial cells on the action of HI-2 we assessed whether reducing both insulin receptors and IGF-1 receptors also reduces hybrid receptors in HUVEC. siRNAs targeting insulin receptor and IGF1 receptor significantly reduced their protein levels (**Figure 4A, left and right**, respectively) compared to controls. Co-transfection with both insulin and IGF1 receptor siRNA decreased hybrid expression in HUVECs compared to control levels (**Figure 4B**). Consistent with HI-2 effect being specific to its effect on hybrids reducing Hybrids using siRNA blunted the effect of HI-2 on ^Ser473^pAkt (**Figure 4C**).

**Figure 4.**
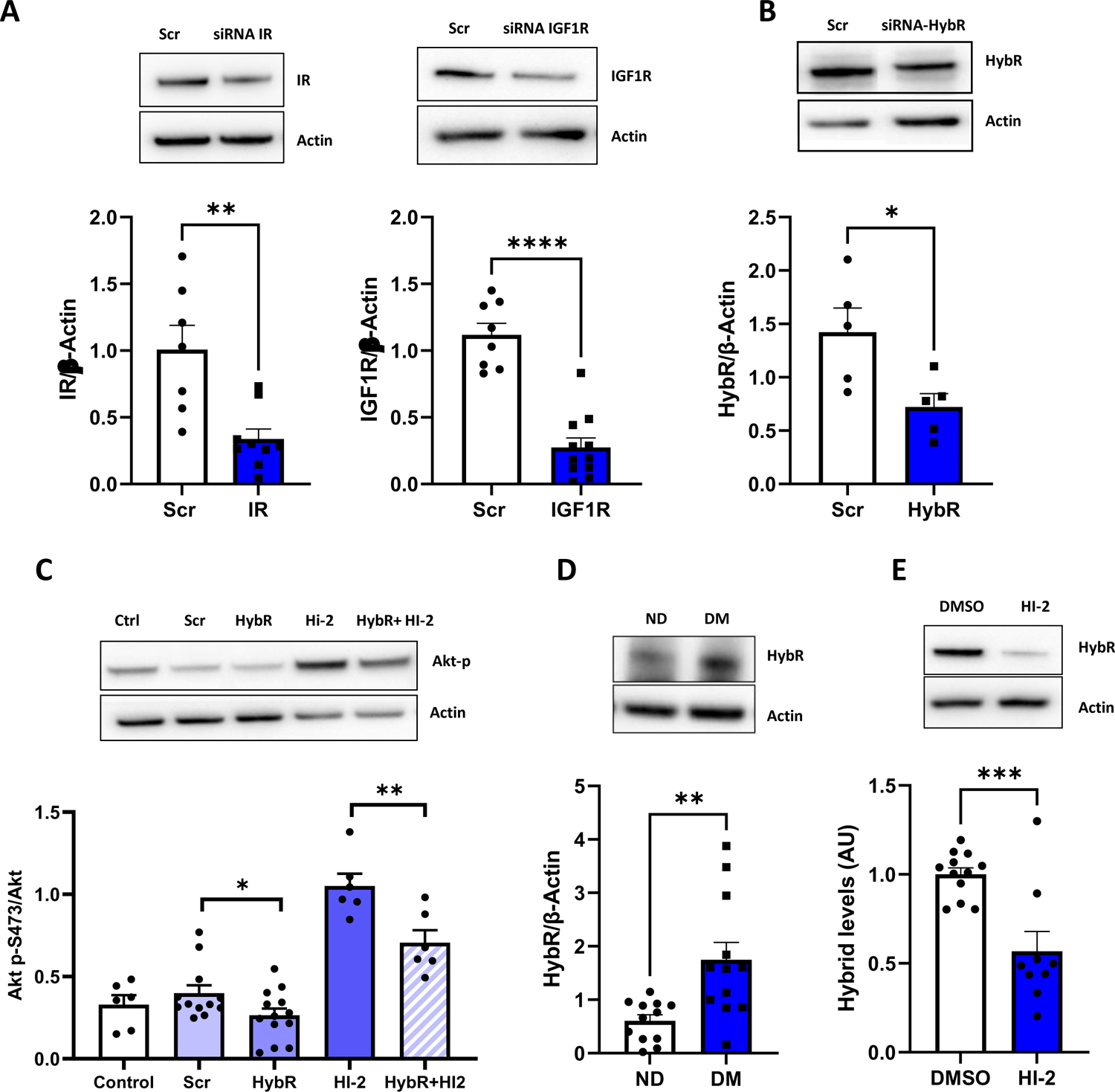
Effect of HI-2 phosphorylation of Akt in endothelial cells and expression of hybrid receptor in primary endothelial cells from patients with advanced coronary artery disease and type 2 diabetes. A) Expression of IR and IGF1R knockdown in HUVECs treated with appropriate siRNA (siRNA IR or siRNA-IGF1R) versus a scrambled siRNA, Scr (n=5, p= 0.0024 for IR and <0.0001 for IGF1R). B) Reduction in hybrid receptor expression in HUVECs treated with appropriate siRNA (siRNA-HybR) versus scrambled siRNA, Scr (n=5, p= 0.0271). C) Ser473 pAkt protein in HUVECs treated with siRNA (siRNA HybR) and HI-2 compared to siRNA alone to reduce hybrid expression levels (100 µM, 24 h treatment, n= >5, p= 0.0091). D) Expression of hybrid receptor levels (HybR) in saphenous vein endothelial cells (SVEC) in diabetic (DM) versus non-diabetic (ND) patient samples. To estimate the relative levels of hybrids, cell lysates were immunoprecipitated with anti-IGF1R antibody, and the levels of IR and IGF1R in immunoprecipitation, supernatant and cell lysate before immunoprecipitation determined. The IR in immunoprecipitation represented IR from hybrids, whereas IR remained in supernatant were likely homo-IR/IR. (n=12 p=0.0032). E) Hybrid receptor protein levels in saphenous vein endothelial cells (SVEC) from patients with diabetes, treated with HI-2 for 24h compared to vehicle control, DMSO. To estimate the relative levels of hybrids, cell lysates were immunoprecipitated with anti-IGF1R antibody, and the levels of IR and IGF1R in immunoprecipitation, supernatant, and cell lysate before immunoprecipitation determined. The IR in immunoprecipitation represented IR from hybrids, whereas IR remained in supernatant were likely homo-IR/IR. (HI-2; 100 µM, 24h treatment, n=9, p=0.0005). All values are expressed as mean and standard error of the mean (SEM). Statistical significance was determined by the Student’s t-test. ns denotes non-significant differences found. DMSO is vehicle control.

### 3.7 Effects of HI-2 on primary endothelial cells from patients with advanced coronary artery disease and type 2 diabetes

To enhance the translational relevance of our findings, we quantified hybrid expression in isolated primary saphenous vein endothelial cells (SVEC) from patients undergoing coronary artery bypass surgery. For the first time, we show that type 2 diabetes is associated with an increase abundance of hybrids in EC from these patients (**Figure 4D**). We then examined the effect of HI-2 on hybrid levels in SVEC from patients with advanced coronary artery disease and type 2 diabetes. HI-2 effectively reduced hybrid protein expression by >50% in SVECs (**Figure 4E**). These data suggest that depletion of hybrid receptor suffices to decrease hybrid-mediated detrimental effects in diabetes, and thus potentially improve ^Ser473^pAkt-mediated metabolic effects in endothelial cells.

### 3.8 Absorption, distribution, metabolism, and excretion analysis

Early-stage physicochemical profiling of HI-2 and analogues to ascertain compound metabolic stability and aqueous solubility *in vitro* (**Table S4**) showed that the HI series has a short half-life in both human and murine liver microsomes. Chemical safety assessment of HI-2 using Derek [45] and Sarah Nexus software [46] suggested the scaffold possessed no obvious safety issues.

## 4. Discussion

Here, we describe the rational, structure-based development of a first-in-class quinoline-containing heterocyclic small molecule that inhibits formation of insulin receptor: IGF-1 receptor heterodimers (hybrid receptors). Using this tool compound, we show for the first time: **1)** That it is technically possible to inhibit hybrid formation in endothelial cells while maintaining expression of the individual receptors. **2)** Reducing hybrid formation in human umbilical vein endothelial cells does not affect cell survival. **3)** Reducing hybrid formation in human umbilical vein endothelial cells reduces expression of the inhibitory p85α subunit of PI3-K signalling. **4)** Consistent with this, reducing hybrid formation enhances basal serine/threonine phosphorylation of Akt. **5)** Reducing hybrids in HUVECs enhances insulin *and* IGF-1 stimulated phosphorylation of Akt, thus enhances Akt activity. **6)** Primary saphenous vein endothelial cells from patients with type 2 diabetes mellitus have increased hybrids, which are reduced after treatment with our tool compound, HI-2.

### A tool compound to inhibit hybrid formation

A pathophysiological hallmark of obesity and type 2 diabetes mellitus is the disruption of insulin signalling in its target cells including vascular endothelial cells [47]. Insulin binds to its cognate receptor and its homologue IGF-1R to stimulate a signalling cascade, proximal in which are the critical signalling nodes phosphatidylinositol 3-kinase (PI3-K) and its downstream target Akt.

IR and IGF-1R are homodimers, which can heterodimerize to form hybrid receptors [48], composed of one haploreceptor of IR and one of IGF-1R. Our group and many others have exploited *in vivo* and *in vitro* genetic approaches to dissect the role of IR [39, 49] and IGF-1R [19, 50] in endothelial cell signalling (for review see [51]). The lack of amenability of hybrids to genetic manipulation makes examining their role in endothelial cell physiology challenging. Preventing the protein-protein interaction between IRαβ and IGF1Rαβ hybrids, using a small molecule is one approach with a potential to circumvent this problem [48]. We set out to design, synthesise and test small molecules that selectively inhibit hybrid formation. As there is no published structure for hybrids, we generated a hybrid homology model based on the IR ectodomain dimer [29] a valid structure to employ due to the high homology between IR and IGF-1R. We used this model to identify potentially important regions involved in hybrid formation. A region covering amino acids 400-570 was chosen for virtual high-throughput screening as from three hotspots identified, this region had the lowest degree of sequence identity between IR and IGF-1R, affording us the greatest likelihood of identifying a selective inhibitor of hybrid formation. Virtual high throughput screening identified compounds predicted to bind to IR and prevent hybrid formation. A range of molecules were screened for compound purity and integrity before a LIVE-DEAD assay ascertained compound toxicity in HEK293 cells. To examine the effect of these molecules on hybrid formation, we developed a Bioluminescence Resonance Energy Transfer (BRET) assay [37] as our initial screen, before moving on to test our most promising compound in a range of *in vitro* assays using endothelial cells.

### Effect of reducing hybrids on endothelial cell insulin signalling

We examined the effect of HI-2 in EC using HUVECs. When HUVECs were treated with HI-2, hybrid receptor levels were reduced with no effect on total IGF-1R or IR expression demonstrating the feasibility of a small molecule approach to inhibiting hybrid formation. Downstream of the IR/IGF-1R/hybrid signalling node is the phosphoinositide-3-kinase/Akt pathway [52]. Under unstimulated conditions, HI-2 reduced the expression of the negative p85α regulatory subunit of phosphoinositide-3-kinase. Consistent with this, HI-2 also induced an increase in basal serine/threonine phosphorylation of Akt raising the possibility that hybrids have a selective tonic inhibitory effect on PI3-K activity, not seen when reducing IR or IGF-1R expression in endothelial cells.

To examine this further, we investigated the effect of HI-2 on insulin and IGF-1 stimulated activation of the phosphoinositide-3-kinase/Akt pathway. Following stimulation with insulin, autophosphorylation of tyrosine residues on the β subunit of IR/IGF1R initiates phosphorylation of IR substrate (IRS) proteins at multiple tyrosine residues [53]. Phosphorylated IRS1 subsequently binds phosphatidylinositol 3-kinase, initiating the conversion of the plasma lipid phosphatidylinositol 3,4,-bisphosphate to phosphatidylinositol 3,4,5-trisphosphate (PIP3). Phosphoinositide-3-kinase is a heterodimer consisting of one catalytic subunit (p110α, p110β, or p110δ) and one regulatory subunit (p85α, p85β, p85γ, p50α or p55α). The p110α isoform is ubiquitously expressed and is essential for proliferation during growth-factor signalling, and oncogenic transformation [54]. Accumulating evidence suggests that changes in p85α levels modulate phosphoinositide-3-kinase activation. It has been suggested that monomeric free p85α may act as a negative regulator of Phosphoinositide-3-kinase signalling [55]. Phosphoinositide-3-kinase activates the downstream serine−threonine kinase Akt [56], for the full activation of Akt, phosphorylation of a serine and threonine residue is required. Consistent with our findings in unstimulated HUVECs, we showed that HUVECs treated with HI-2 had enhanced Akt (serine/threonine) phosphorylation in response to insulin and IGF-1. Moreover, the phosphoinositide-3-kinase inhibitor, wortmannin blunted the effect of HI-2 on Akt phosphorylation, suggesting that the HI-2-stimulated Akt phosphorylation is mediated via the PI3-K signalling pathway.

### Effect of reducing hybrids on endothelial cell responses to shear stress

In endothelial cells, it has been shown that shear stress increases phosphorylation of Akt in a PI3-K dependent, but receptor tyrosine kinase independent fashion [57]. We therefore examined the effect of HI-2 on shear stress-induced phosphorylation of Akt. Consistent with our findings, in insulin and IGF-1 response studies, we showed that HI-2 augments shear stress-induced phosphorylation of Akt. These data support the intriguing possibility that hybrid receptors may have a unique role in regulation of the PI3-K/Akt signalling pathway *per se*.

### Reduction in hybrids suggests they have physiological functions distinct to insulin receptor and IGF-1 receptors

Our dataset raises the exciting possibility that hybrids may have a biological signature distinct from their component receptor types IR and IGF-1R. The effect of our hybrid inhibitor does not recapitulate the phenotype of endothelial cells with either reduced IR/IGF-1R [19] or alternative approaches to reducing insulin and IGF-1R signalling at the IR/IGF-1R/hybrid signalling node[58, 59]. Neither is the effect due to reduced IR signalling. Our own data [58] shows that knockdown of the IR in HUVECs, in contrast to the effect seen in reducing hybrids here, led to blunted Akt activation. In the same study, we showed that knockdown of IR in HUVECs leads to reduced angiogenic sprouting, whereas knockdown of hybrid receptors increased angiogenic sprouting. Data regarding deletion of IGF-1R in the endothelium also support a discrete role for hybrids. IGF-1 has been shown to increase phosphorylation of Akt in endothelial cells and enhance angiogenesis [60]. In another study, providing supporting data from a separate cell type macrophages deficient in IGF-1R had blunted IGF-1 mediated Akt phosphorylation, with no effect on insulin-induced phosphorylation of Akt [61].

To examine the possibility that increased endothelial hybrid expression is a feature type 2 diabetes in humans, we measured hybrid expression in endothelial cells from saphenous vein isolated from patients with advanced coronary artery disease. We confirmed that hybrid expression was significantly higher in saphenous vein endothelial cells from patients with type 2 diabetes mellitus. Additionally, in terms of translational relevance and as a foundation for future investigations extending beyond the scope of this report, we demonstrated that our hybrid inhibitor successfully decreased hybrid receptors in patient samples.

### Study limitations

To progress this series towards *in vivo* analysis of hybrid function, new analogues of HI-2 would need to possess improved metabolic stability and aqueous solubility. Compound aqueous solubility was measured at pH 7.4 in phosphate buffered saline (PBS) using the method described by Bellenie et. Al.[62] Our initial hit, HI-1 possessed moderate solubility, but solubility was improved by replacement of the carbo-aromatic rings with heterocycles, correlating well with cLogP values for the series. Our current series of inhibitors lack the potency and physicochemical properties to pursue an *in vivo* series of experiments, to understand fully the role of hybrid receptors in a murine model of metabolic disease. Given the recently published structure of the hybrid receptors with IGF-1 bound[63], future work will seek to address issues with potency and physicochemical properties in a structure-guided manner.

## Conclusions

We set out to develop a tool compound that inhibits hybrid formation in human cells. Here, we present a novel small molecule that selectively inhibits the formation of hybrid receptors in human EC. We have shown that reducing hybrid expression in HUVECs alters expression and/or activation status of molecules at two key nodes in the insulin signalling pathway, including the p85α subunit of phosphatidylinositol 3-kinase (PI3-K) and Akt. Moreover, we show that this tool compound enhances serine phosphorylation of Akt, in response to insulin and mechanical shear stress. Development and refinement of small molecules targeting hybrid receptors could pave the way for innovative therapeutic approaches against insulin resistance and, potentially, cardiovascular complications.

### Author contributions

MK and KJS designed and directed the study. KJS carried out the molecular modelling and compound selection. DJB and CWGF contributed to the study design. CGM, HV, NJH and KB carried out and analysed western blotting data. KJS, ST, LL and MJM synthesised HI analogues. TI provided IR-Rluc and IGF1R-YPET plasmids for BRET assays. All authors, including EC, PM, SBW and RC analysed the data and discussed the results. MK, KJS and CW prepared the manuscript. Prof Mark Kearney is the guarantor of this work and, as such, had full access to all the data in the study and takes responsibility for the integrity of the data and the accuracy of the data analysis.

## Acknowledgements

The authors acknowledge Dr Christopher M. Pask for small molecule X-ray crystallography experiments and Ms Khanh Pham (LICAMM-University of Huddersfield Placement Programme) for her technical support. The authors would like to thank Dr Karen Porter for the donation of saphenous vein endothelial cells.

Appendix A. Supplementary data

## Data Availability

Data will be available on request.

