## Supplementary data for "Small molecule modulation of insulin receptor-insulin like growth factor-1 receptor heterodimers in human endothelial cells"

<u>**Supplementary Material**</u>

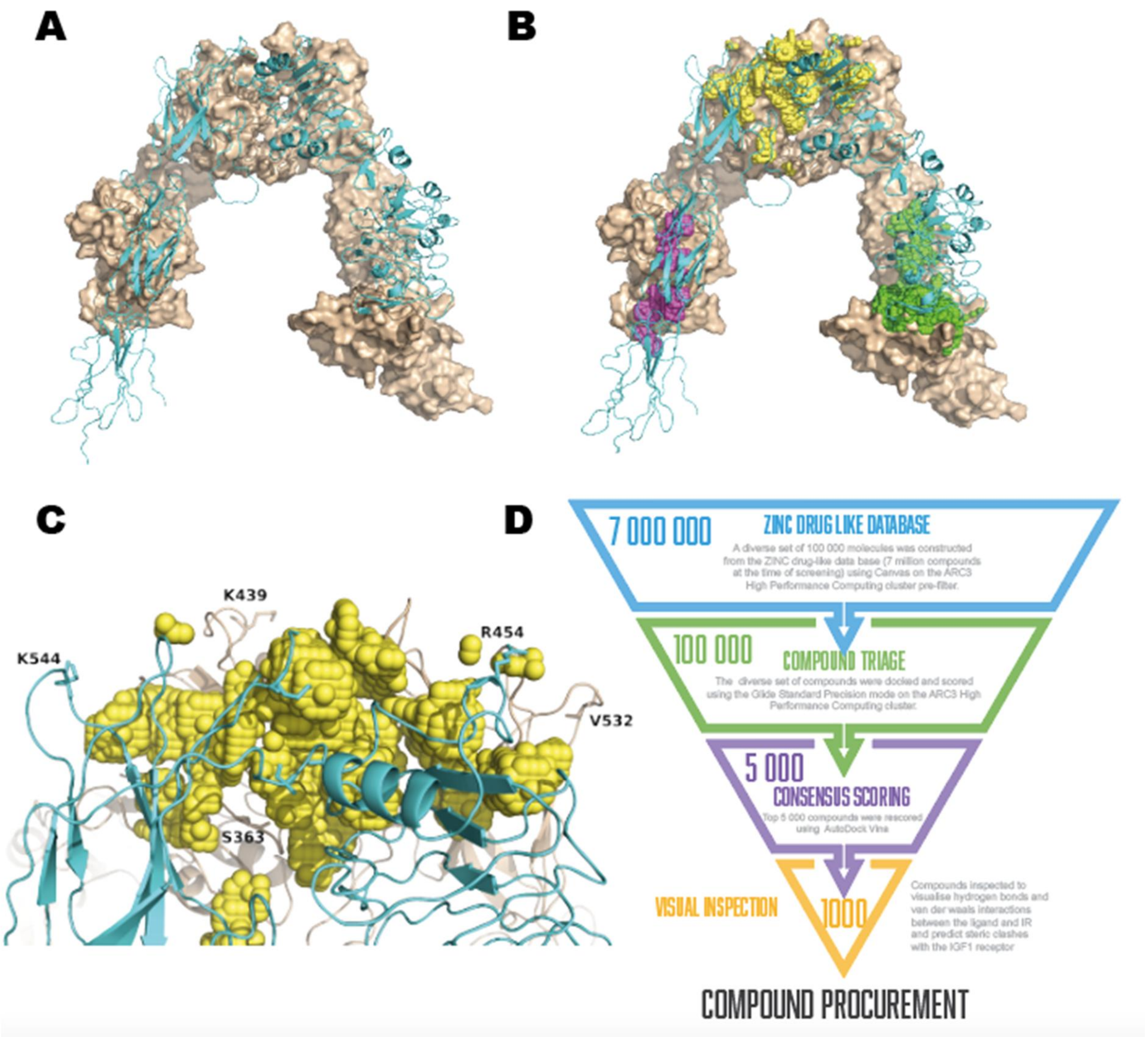


**Figure S1: A-** Homology model of the IR-IGF-1R hybrid receptor produced using the I-TASSER webserver and the IR ectodomain dimer structure (PDB ID 4ZXB). The IR monomer is shown as a wheat surface and the IGF-1R monomer is shown as cyan ribbons. Image generated using Pymol. **B-** Hotspots identified as critical for IR-IGF1R hybrid receptor formation using the KFC2 webserver. The IR monomer is shown as a wheat surface and the IGF1R monomer is shown as cyan ribbons. Hotspot 1 is shown as green spheres, hotspot 2 is shown as yellow spheres and hotspot 3 is shown as lilac spheres. Image generated using Pymol **C-** Hotspot 2 (yellow spheres) covers residues 388-557 in the IR (wheat ribbons) and residues 398-576 in the IGF1R (cyans ribbons) A number of residues involved in the interaction are highlighted as wheat/cyan sticks. Image generated using Pymol. **D-** Overview of the virtual high-throughput screening cascade using to identify potential inhibitors of hybrid formation. A diverse set of 100 000 molecules was constructed from the ZINC drug-like data base, the top scoring compounds using the Glide SP scoring function were then rescored using the AutoDock Vina scoring function, before the top 1000 compounds were visually inspected to visualise hydrogen bonds and van der waals interactions between the ligand and IR and predicted steric clashes with the IGF-1R

**Table S1**: Antibodies used in this paper

| Antibody Name | Antibody ID | Manufacturer | Source/Isotype | Concentration |
|---|---|---|---|---|
| Insulin Receptor β (4B8) | #3025 | CST | Rabbit | 1:1000 |
| IGF1R β (D23H3) XP® | #9750 | CST | Rabbit | 1:1000 |
| β-actin (C4) | #sc-47778 | SCBT | Mouse | 1:5000 |
| Akt pan (11E7) | #4685 | CST | Rabbit | 1:1000 |
| Phospho-Akt (Ser473) (D9E) XP® | #4060 | CST | Rabbit | 1:1000 |
| Phospho-Akt (Thr308) (244F9) | #4056 | CST | Rabbit | 1:1000 |
| PI3 Kinase p85 (19H8) | #4257 | CST | Rabbit | 1:1000 |
| PI3 Kinase p110α (C73F8) | #4249 | CST | Rabbit | 1:1000 |
| PI3 Kinase p110β (C33D4) | #3011 | CST | Rabbit | 1:1000 |
| Rabbit IgG HRP Linked Whole Ab | #NA9341ML | SLS | Anti-Rabbit HRP | 1:10 000 |
| Rabbit anti-Mouse IgG (H+L), Superclonal Recombinant Secondary Antibody, HRP | #A27025 | ThermoFisher | Anti-Mouse HRP | 1:10 000 |

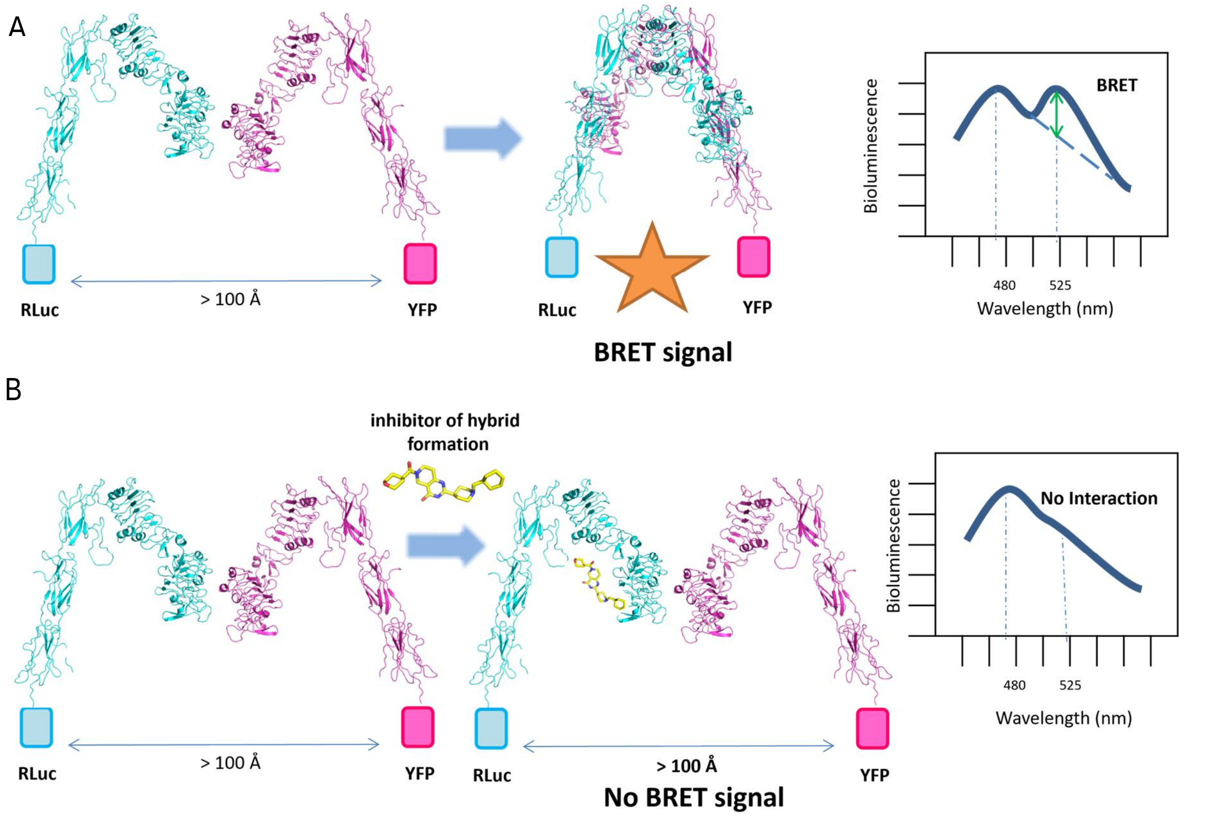


**Figure S2**: Overview of the BRET assay A) To study the interaction between two proteins, IR is fused to Renilla luciferase (Rluc) and IGF1R is fused to a yellow fluorescent protein (YPET). The reaction is initiated by addition of the substrate of luciferase, coelenterazine. If the distance between IR and IGF1R is 10 to 100 Å, part of the energy of the excited Rluc is transferred to the YPET, resulting in an additional signal emitted by the YFP. B) If hybrid formation between IR and IGF1R is inhibited by a small molecule, light is emitted with an emission spectrum characteristic of the Rluc only.

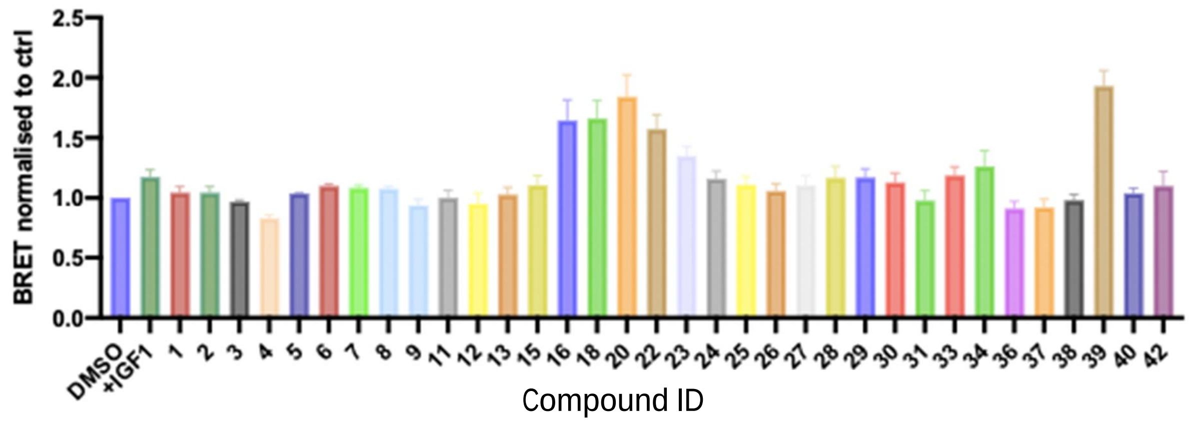


**Figure S3**: Compounds identified using vHTS and screened using the BRET assay. Compound structures are displayed in Table S3. All compounds were screened at 100 µM final concentration (n=8,3).

**Table S2: Compounds selected from virtual high-throughput screening.**

| | Cpd ID | Structure | Source |
|---|---|---|---|
| **1** | **5373400** | 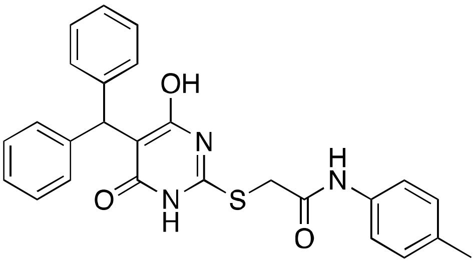 | Chembridge |
| **2** | **6520773** | 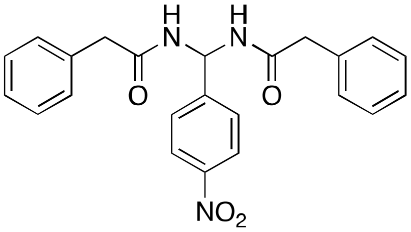 | Chembridge |
| **3** | **7911669** | 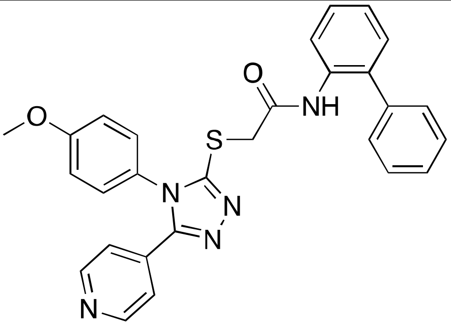 | Chembridge |
| **4** | **7922787** | 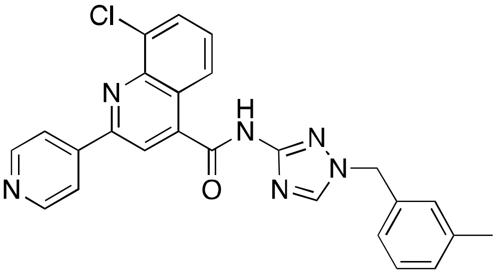 | Chembridge |
| **5** | **16312697** | 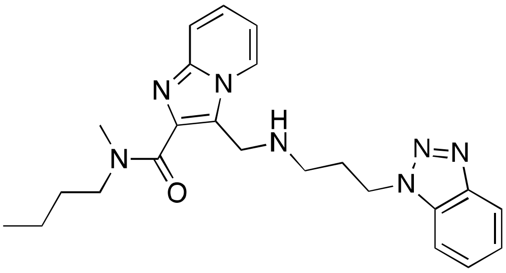 | Chembridge |
| **6** | **39752577** | 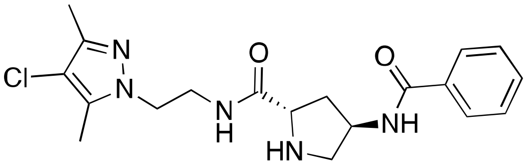 | Chembridge |

| | | | |
|---|---|---|---|
| **7** | **68092032** | 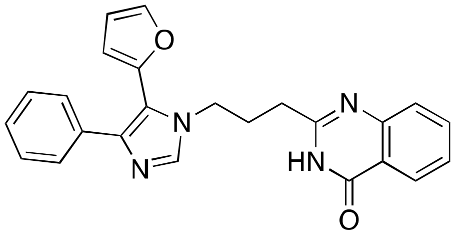 | Chembridge |
| **8** | **88193220** | 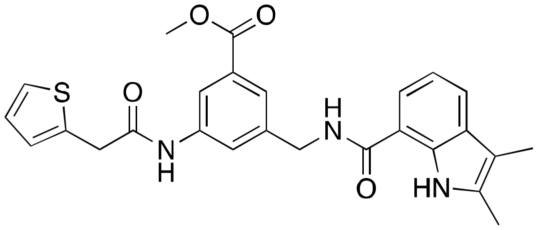 | Chembridge |
| **9** | **27657671** | 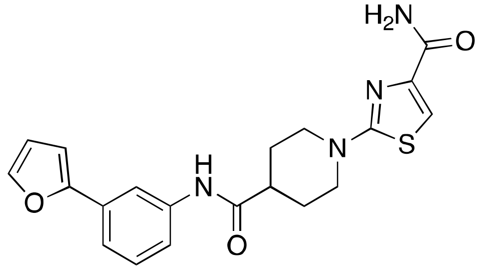 | Chembridge |
| **10** | **28571425** | 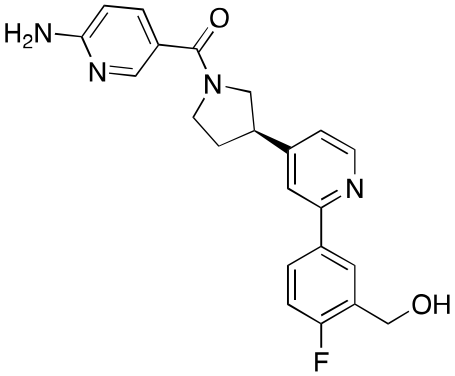 | Chembridge |
| **11** | **51921735** | 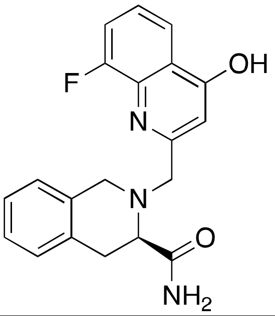 | Chembridge |
| **12** | **9108461** | 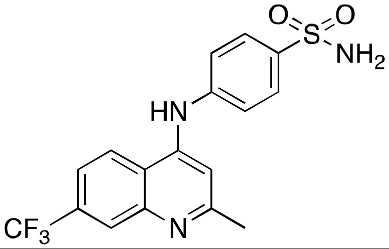 | Chembridge |

| | | | |
|---|---|---|---|
| **13** | **27150262** | 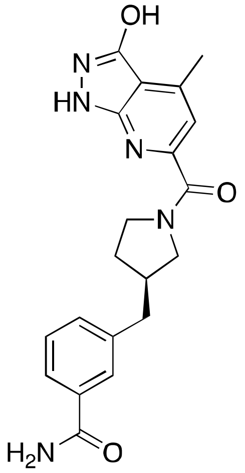 | Chembridge |
| **14** | **91410900** | 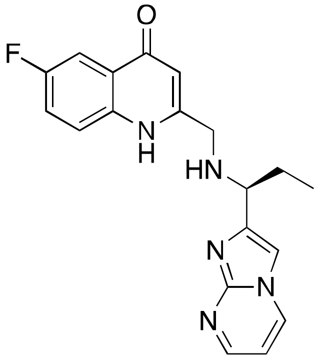 | Chembridge |
| **15** | **66215718** | 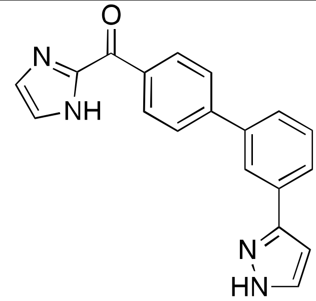 | Chembridge |
| **16** | **10907689** | 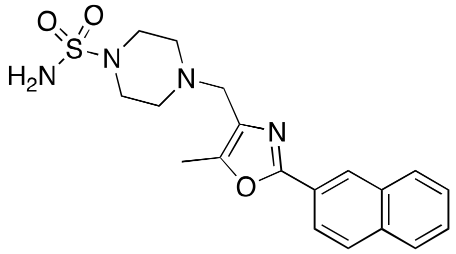 | Chembridge |
| **17** | **87179763** | 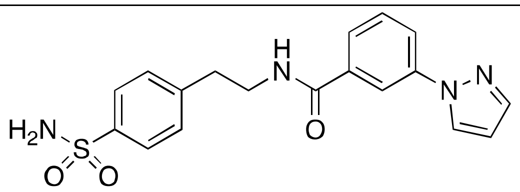 | Chembridge |
| **18** | **5908526** | 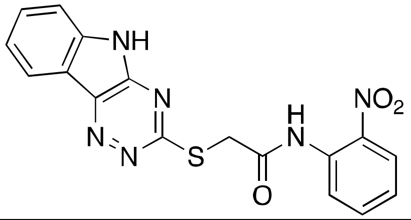 | Chembridge |

| | | | |
|---|---|---|---|
| **19** | **45256902** | 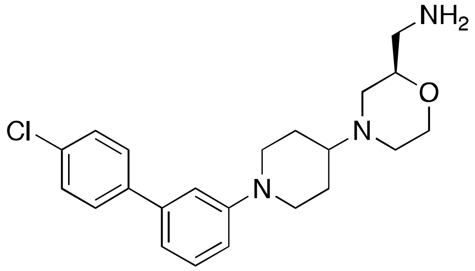 | Chembridge |
| **20** | **7959117** | 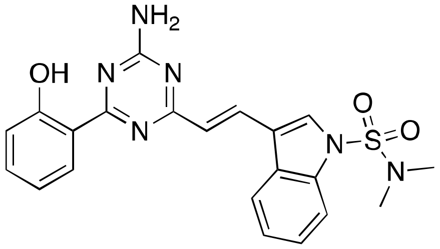 | Chembridge |
| **21** | **61771203** | 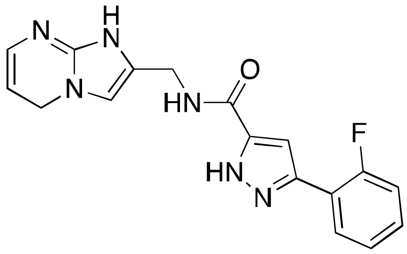 | Chembridge |
| **22** | **9265357** | 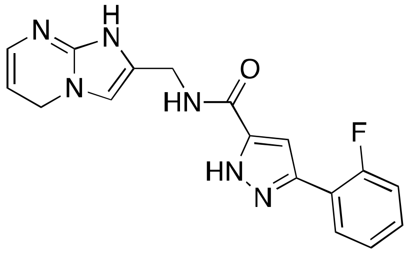 | Chembridge |
| **23** | **56945281** | 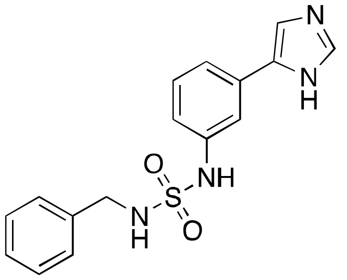 | Chembridge |
| **24** | **97997697** | 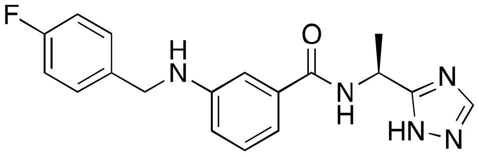 | Chembridge |
| **25** | **56206121** | 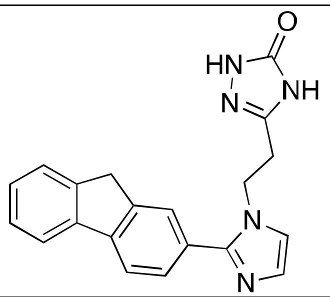 | Chembridge |

| | | | |
|---|---|---|---|
| **26** | **9256200** | 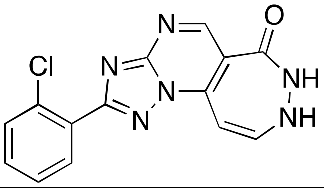 | Chembridge |
| **27** | **58962706** | 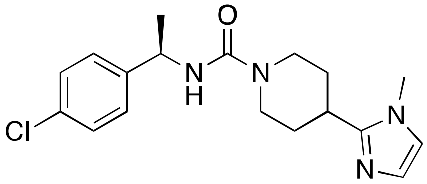 | Chembridge |
| **28** | **36106297** | 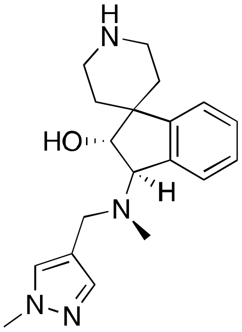 | Chembridge |
| **29** | **78321474** | 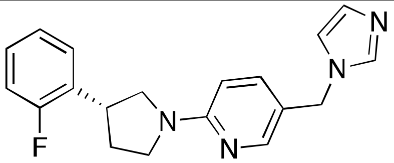 | Chembridge |
| **30** | **5580755** | 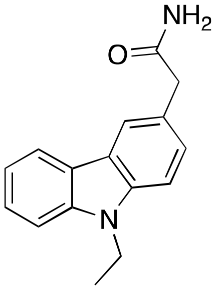 | Chembridge |
| **31** | **71511441** | 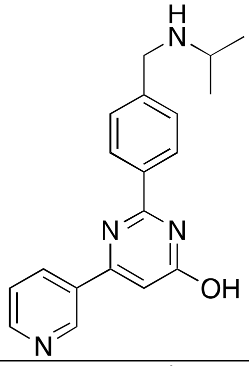 | Chembridge |
| **32** | **20178901** | 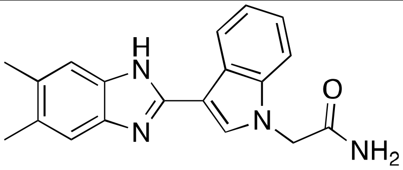 | Chembridge |

| | | | |
|---|---|---|---|
| **33** | **RJC00584** | 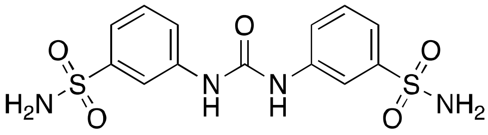 | Maybridge |
| **34** | **BTB13888** | 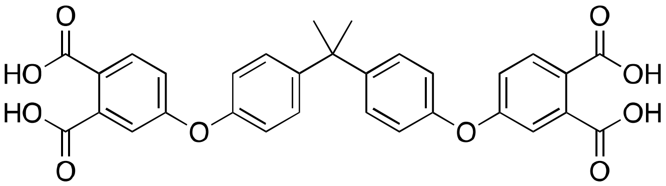 | Maybridge |
| **35** | **GK00878** | 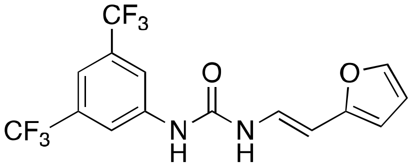 | Maybridge |
| **36** | **HTS03564** | 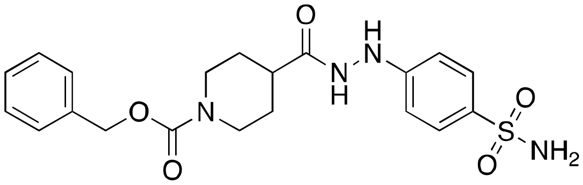 | Maybridge |
| **37** | **SCR01167** | 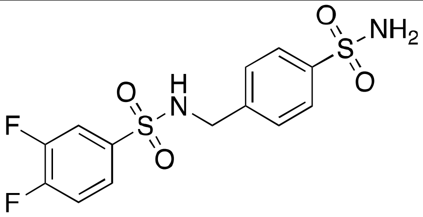 | Maybridge |
| **38** | **SEW06378** | 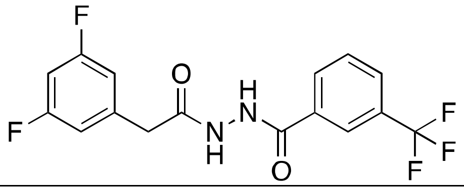 | Maybridge |
| **39** | **BTB01314** | 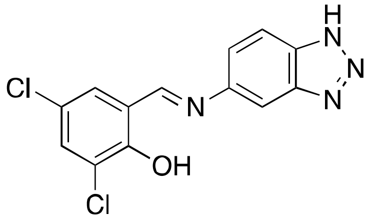 | Maybridge |
| **40** | **BTB14921** | 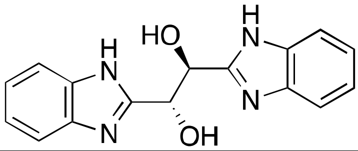 | Maybridge |
| **41** | **MGH00165** | 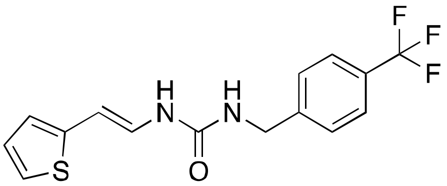 | Maybridge |

| | | | |
|---|---|---|---|
| **42** | **HTS12296** | 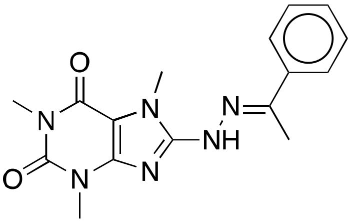 | Maybridge |

**Table S3: Analogues of HI-1 synthesised and purchased**

| | Cpd ID | Structure | Source |
|---|---|---|---|
| **4** | **HI1** | 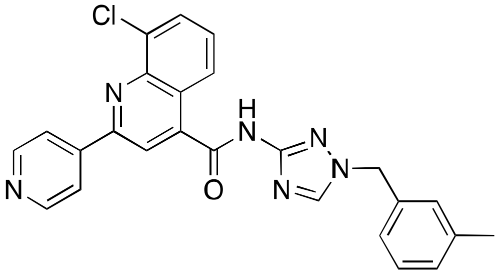 | Chembridge/ in house |
| **43** | **HI2** | 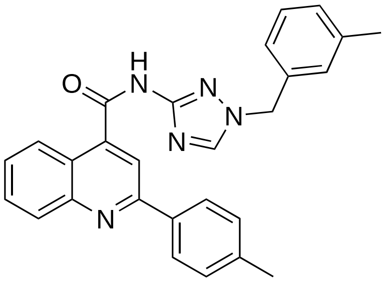 | In-house |
| **44** | **MJM417** | 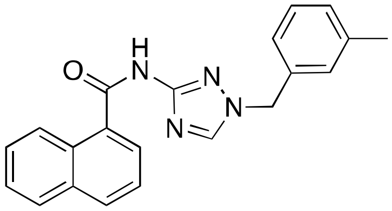 | In-house |
| **45** | **MJM 423** | 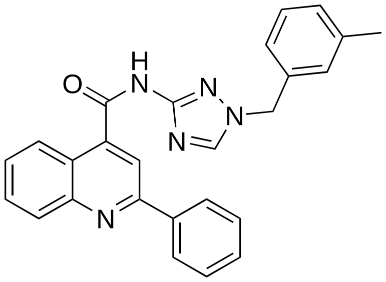 | In-house |
| **46** | **MJM428** | 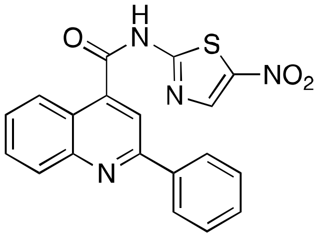 | In-house |

| | | | |
|---|---|---|---|
| **47** | **LL39** | 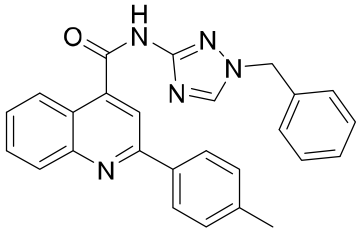 | In-house |
| **48** | **LL40** | 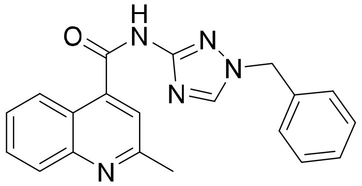 | In-house |
| **49** | **LL49** | 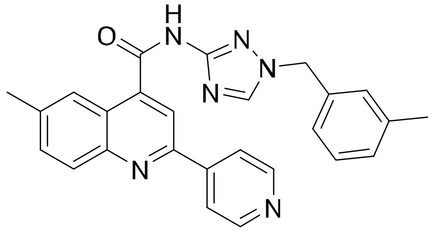 | In-house |
| **50** | **LL55** | 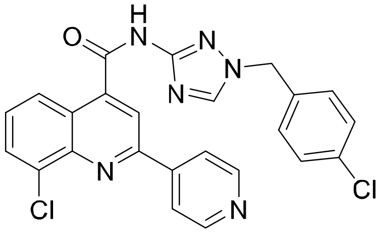 | In-house |
| **51** | **SJT1** | 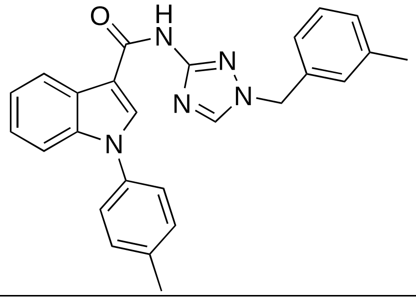 | In-house |
| **52** | **SJT2** | 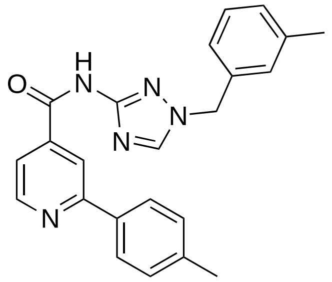 | In-house |
| **53** | **MJM448** | 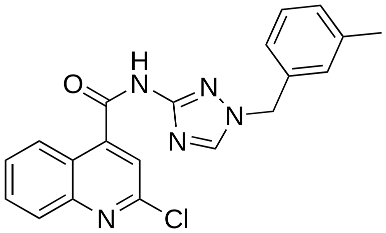 | In-house |

| | | | |
|---|---|---|---|
| **54** | **MJM452** | 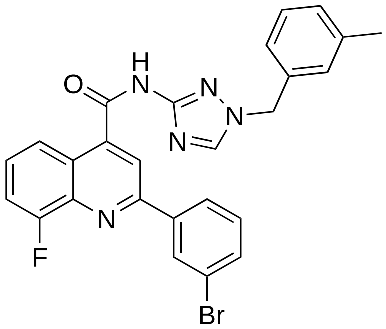 | In-house |
| **55** | **MJM454** | 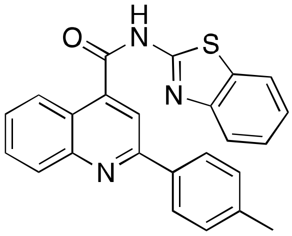 | In-house |
| **56** | **MJM456** | 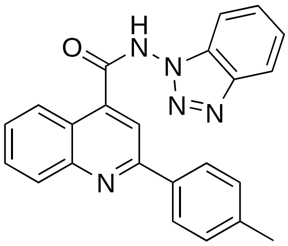 | In-house |
| **57** | **STK461970** | 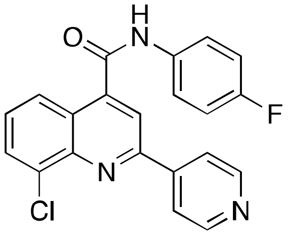 | Vitas-M Labs |
| **58** | **STK465947** | 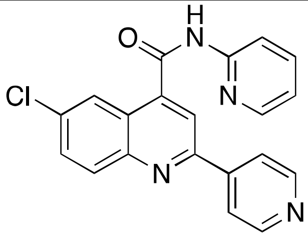 | Vitas-M Labs |
| **59** | **STK465127** | 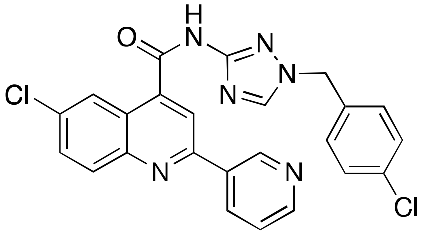 | Vitas-M Labs |
| **60** | **STK969012** | 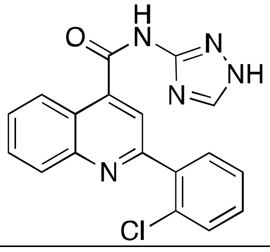 | Vitas-M Labs |

| | | | |
|---|---|---|---|
| **61** | **STK446492** | 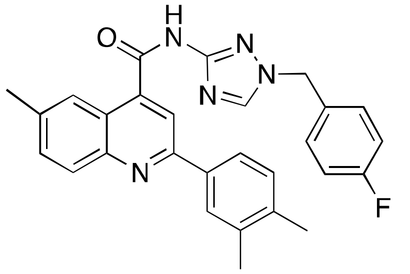 | Vitas-M-Labs |
| **62** | **STK418759** | 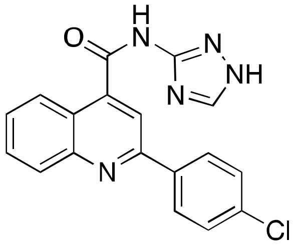 | Vitas-M Labs |
| **63** | **STK464985** | 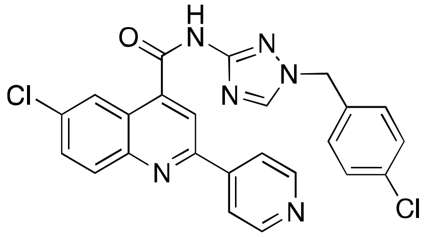 | Vitas-M Labs |
| **64** | **Talnetant** | 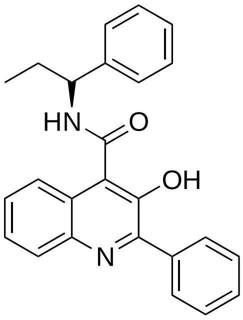 | GlaxoSmithKline |
| **65** | **ALB-H11666337** | 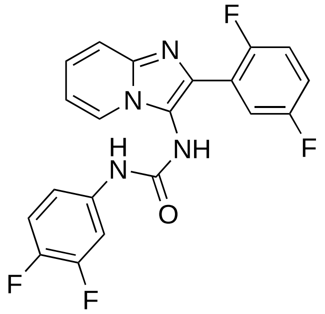 | AMRI |
| **66** | **ALB-H05945621** | 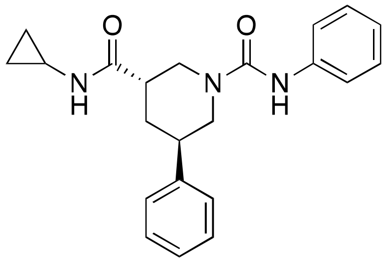 | AMRI |

**Table S4: Solubility and half-life measurements for selected HI analogues**

| | Structure | Solubility (pH 7.4) µM | $T_{1/2}$ (min) | |
|---|---|---|---|---|
| | | | **Human** | **Mouse** |
| **4 (HI1)** | 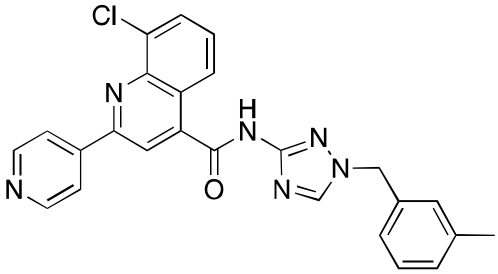 | 18.4 | 5.77 | 5.93 |
| **43 (HI2)** | 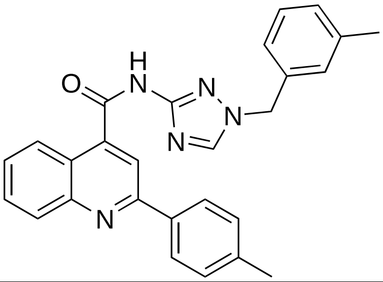 | 1.8 | 6.97 | 1.03 |
| **51** | 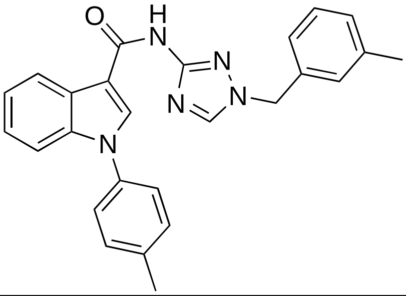 | 1.66 | ND | ND |
| **52** | 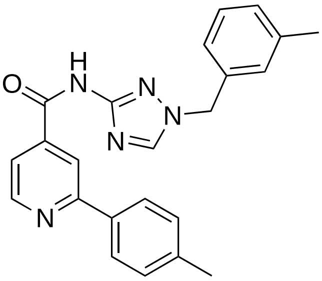 | 12.9 | ND | ND |
| **53** | 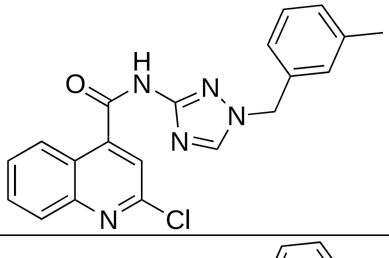 | 75.6 | ND | ND |
| **54** | 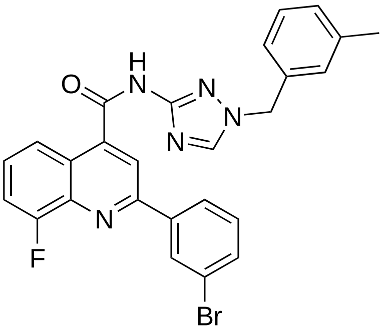 | 0.09 | ND | ND |

| | | | | |
|---|---|---|---|---|
| **55** | 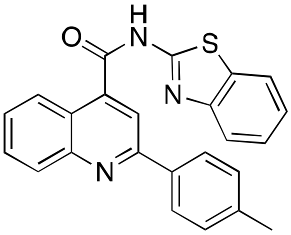 | 0.37 | ND | ND |
| **56** | 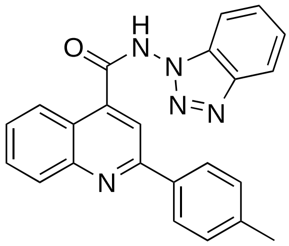 | 87.2 | ND | ND |

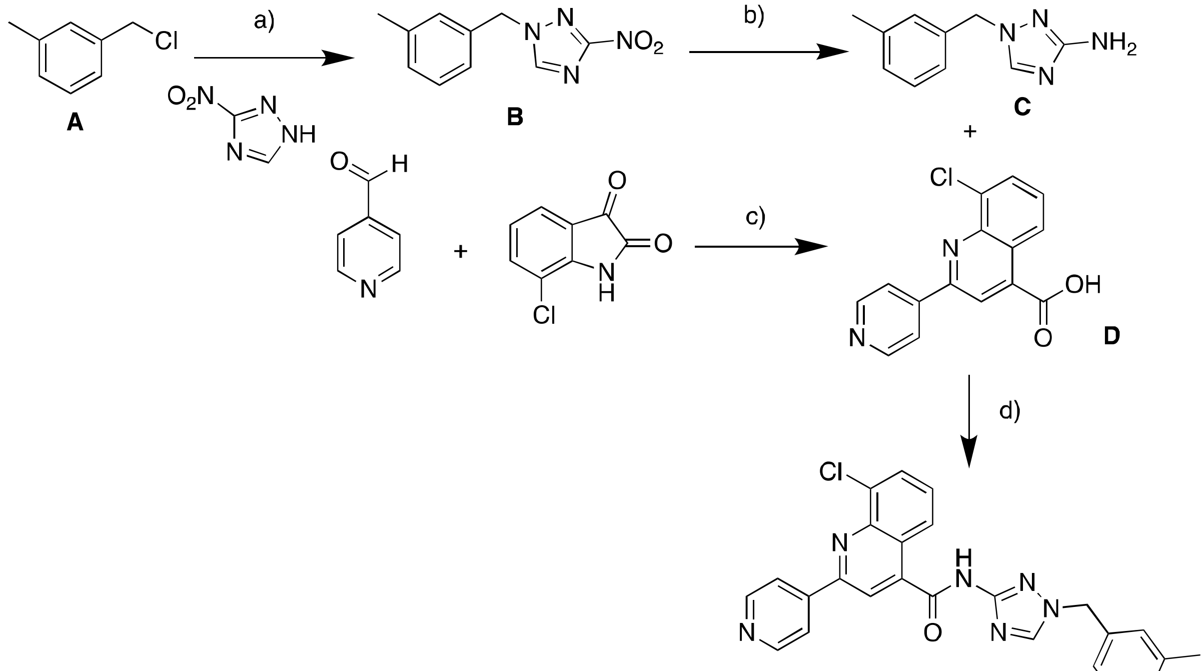

**HI-1**

**Scheme S1**- Example Synthesis of HI1 a) NaOEt, EtOH, 80 °C, 18 h, 99% b) $H_2$, Pd/C, EtOH r.t., 8 h, 95% c) KOH, EtOH aq, 120°C MW, 64% d) $T_3P$, $NEt_3$, EtOAc, 80 °C, 18 h, 7%

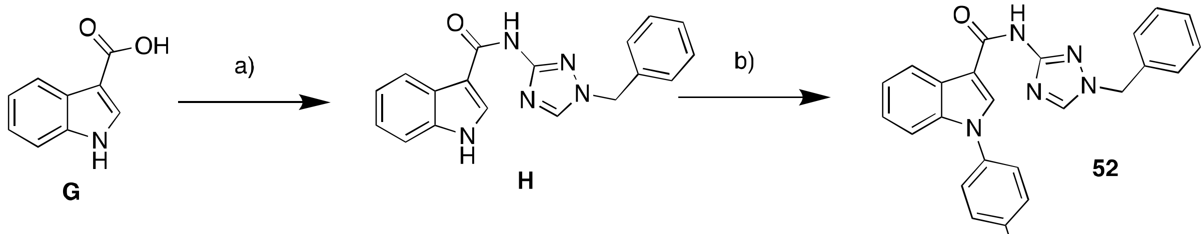

**Scheme S2**- Example Synthesis of Indole Analogue 52 a) 1-(3-methylbenzyl)-3-amino-1H-1,2,4-triazole, $T_3P$, $NEt_3$, EtOAc, 80 °C, 18 h, 59%, b) 4-Bromo-toluene, CuI, N,N′-dimethylethylenediamine, $K_3PO_4$, DMF, 110 °C, 18 h, 17%

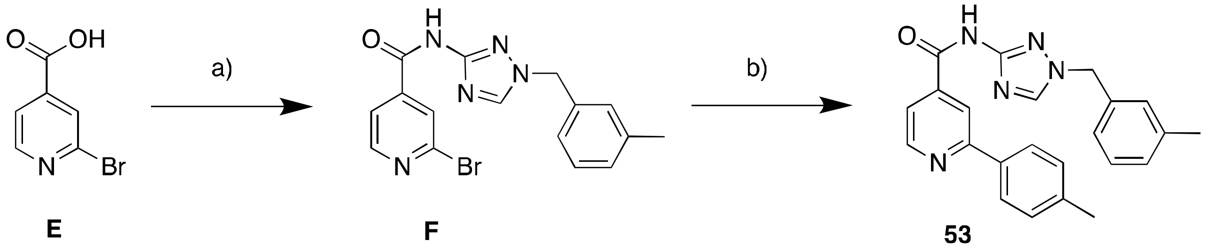


**Scheme S3**- Example Synthesis of Pyridyl Analogue 53 a) 1-(3-methylbenzyl)-3-amino-1H-1,2,4-triazole, $T_3P$, $NEt_3$, EtOAc, 80 °C, 18 h, 59%, b) 4-Tolylboronic acid, $PdCl_2(PPh_3)_2$, $CsCO_3$, toluene, 100 °C, 18 h, 51%.

**Compound Synthesis:** All solvents and reagents were obtained from commercial suppliers and used without further purification. Solvents used were HPLC or analytical grade. Thin layer chromatography was performed on aluminium backed silica gel supplied by Merck, visualised using an ultraviolet lamp. Flash column chromatography was performed using silica gel 60 (40-63μm particles). Automated flash column chromatography was performed on a Biotage® Isolera™ One machine using Biotage® Sfär columns of varying sizes between 5 g and 100 g. Automated reverse phase flash column chromatography was performed using C18 silica columns. Proton and Carbon-13 NMR data were collected on Bruker Avance III 500. All shifts were recorded against an internal standard of tetramethyl silane. Solvents used for NMR (Methanol-$d_4$ and DMSO-$d_6$) were obtained from Sigma-Aldrich. $^1$H NMR data is reported in the following format: ppm (splitting pattern, coupling constant (Hz), number of protons, proton assignment). Signal assignments were deduced with the aid of TopSpin, MestReNova, DEPT 135, COSY, HSQC and HMBC. LC-MS (liquid chromatography-mass spectrometry) data were recorded on a Donex Ultimate 3000 LC system with a MeCN/$H_2O$ +0.1% formic acid gradient. HRMS data were recorded using a Bruker MaXis impact spectrometer using electron spray ionisation. Infrared spectra were recorded on a Perkin-Elmer one FTIR spectrometer. Melting points were recorded on Griffin Education MELTP melting point apparatus. Unless otherwise stated, all reactions were carried out under air and at room temperature. Solvents were removed under reduced pressure using a Büchi rotary evaporator and a Vacuubrand PC2001 Vario diaphragm pump. All other solvents used were of chromatography or analytical grade. Commercially available starting materials were obtained from either Fluorochem, Alfa Aesar or Sigma Aldrich. Flash column chromatography was performed using silica gel 60 (35-70μm particles) obtained from Merck. Thin layer chromatography was performed using pre-coated aluminium plates (Merck silica gel 60 F254) that are commercially available from Merck. An ultraviolet lamp (λmax = 254 nm) and $KMnO_4$ were used for visualisation.

**General Methods**

**Method A-** Anhydrous ethanol (0.5 M) was added to sodium metal (1.1 eq) under inert atmosphere. The resulting mixture was stirred at room temperature until all solids had dissolved. The appropriate triazole (1.0 eq) was added against a counter flow of $N_2$, and the reaction allowed to stir until the solids had dissolved. The appropriate benzyl chloride (1.3 eq.) was added, and the reaction heated to 80 °C until TLC and/or LCMS indicated triazole starting material had been consumed. The reaction was cooled to room temperature, quenched with water (2 mL) and concentrated. The residue was partitioned between water (20 mL), and EtOAc (3 x 20 mL). Organics were combined and washed with brine (20 mL), dried ($Na_2SO_4$), filtered and concentrated to yield the crude product which was purified by column chromatography.

**Method B-** A catalytic amount of activated palladium on charcoal (5-10 mol%) was added to a stirred solution of the appropriate nitro-compound (1.0 eq) in methanol (0.5 M). The mixture was stirred at room temperature under and atmosphere of hydrogen until TLC and/or LCMS indicated nitro starting material had been consumed. The reaction was filtered through celite (washed thoroughly with methanol) and the filtrate concentrated under reduced pressure.

**Method C-** To a solution of the appropriate isatin (1.0 eq) in EtOH/ water (1:1, 5 ml) was added 4-acetylpyridine (1.0 eq) and KOH (5.0 eq). The resulting reaction mixture was stirred until homogenous then heated in a microwave reactor at 120 °C for 10 min. This was cooled, diluted with water (50 ml) and acidified using 2M HCl to pH 6.5. The resulting precipitate was collected by filtration, washed with water (50 ml) and EtOAc (50 ml) and dried in a desiccator to give the crude product which was triturated with hot acetone to give the title compound.

**Method D-** The appropriate carboxylic acid (1.0 eq), $NEt_3$ (6.0 eq) and $T_3P$ (50 % w/w in EtOAc, 4.0 eq) were added to EtOAc (0.5 M) and stirred at room temperature for 10 mins until all solids had dissolved. The appropriate amine (0.9 eq) was added and the reaction heated to 80 °C until TLC and/or LCMS indicated amine starting material had been consumed. The reactions was cooled to room temperature and water added (20 mL). The organics were separated and washed with water (20 mL), saturated $NaHCO_3$ solution (10 mL) and brine (20 mL), dried ($MgSO_4$), filtered and concentrated to yield the crude product which was purified by column chromatography.

**Method E-** The appropriate halide (1.0 eq) and boronic acid (1.2 eq) were dissolved in dioxane (2.3 mL). The solution was purged with $N_2$, $PdCl_2(PPh_3)_2$ (0.1 eq) and $CsCO_3$ (1.0 eq) added. The reaction was purged with further $N_2$ and stirred at 100 °C until TLC and/or LCMS indicated halide starting material had been consumed. The reaction was cooled to room temperature, filtered through celite, and filtrate concentrated. The residue was dissolved in DCM (20 mL), and washed with water (20 mL), sat. aqueous $NH_4Cl$ (20 mL), dried ($Na_2SO_4$) and concentrated to yield the crude product which was purified by column chromatography.

**1-[(3-Methylphenyl)methyl]-3-nitro-1H-1,2,4-triazole**

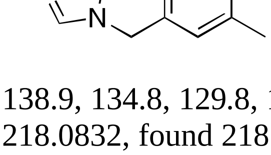


Prepared using Method A. Purification using column chromatography (20-50% EtOAc in Pet. Ether) gave the title compound as a colourless oil. (3.25 g, 14.9 mmol, 99 %) **Rf** 0.20 (33 % petrol in DCM) **$^1$H NMR** (500 MHz, $CDCl_3$) δ = 8.06 (1H, s), 7.31 (1H, t, J = 8.0 Hz), 7.23 (1H, d, J = 8.0 Hz), 7.15 (1H, s), 7.07 (1H, d, J = 8.0 Hz), 5.38 (2H, s), 2.36 (3H, s) **$^{13}$C NMR** (125 MHz, $CDCl_3$) δ = 163.9, 142.5, 138.9, 134.8, 129.8, 128.9, 128.7, 125.0, 53.3, 21.3; **HRMS** (ESI-TOF) *m/z* $[M+H]^+$ calcd for $C_{10}H_{10}N_4O_2$ 218.0832, found 218.0839. **LCMS** RT = 0.60 min, *m/z* = 217.6 $[M+H]^+$.

**1-(4-chlorobenzyl)-3-nitro-1H-1,2,4-triazole**

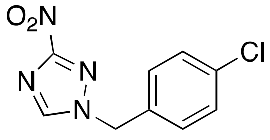


Prepared using Method A. Purification using column chromatography, (20-100% EtOAc in Pet. Ether) gave the title compound (460 mg, 1.93 mmol, 44 %) as a colourless solid. **Rf** 0.0 (5:1 Petrol-EtOAc) **$^1$H NMR** (400 MHz, $CDCl_3$) $\delta$ 8.16 (1H, s), 7.40 (2H, d, J= 8.0 Hz), 7.31 (2H, d, J= 8.0 Hz), 5.42 (2H, s); **$^{13}$C NMR** (400 MHz, $CDCl_3$) $\delta$= 144.6, 135.7, 130.9, 130.0, 129.9, 54.6, 53.5; **HRMS (ESI$^+$)** *m/z*: $[M + H]^+$ Calcd for $C_9H_8N_4O_2$ 239.0336; Found 239.0357. **HPLC** RT = 2.68 min, 100 % relative area.

**1-benzyl-3-nitro-1H-1,2,4-triazole**

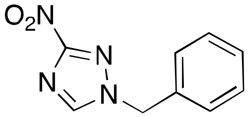


Prepared using Method A. Purification using column chromatography, (20-100% EtOAc in Pet. Ether) gave the title compound (392 mg, 1.92 mmol, 44 %) as a clear oil which solidified on manipulation. **Rf** 0.24 (10 % petrol in DCM) **$^1$H NMR** (400 MHz, $CDCl_3$) $\delta$= 8.14 (1H, s), 7.46-7.40 (3H, m), 7.35-7.29 (2H, m), 5.45 (2H, s); **$^{13}$C NMR** (100 MHz, $CDCl_3$) $\delta$= 162.8, 152.7, 144.7, 132.4, 129. 6, 129.5, 129.0, 128.6, 127.8, 55.5. **HRMS** ($ESI^+$) *m/z*: $[M + Na]^+$ Calcd for $C_9H_8N_4O_2Na$ 204.0647; Found 227.0807. **HPLC** RT = 2.21 min, 100 % relative area.

**1-[(3-Methylphenyl)methyl]-1H-1,2,4-triazol-3-amine.**

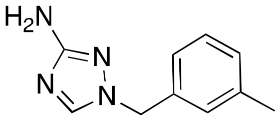


Prepared using Method B to give the title compound as a colourless solid (2.45 g, 13.0 mmol, 95%) which was used without further purification. **Rf** 0.88 (5% MeOH in DCM) **$^1$ H NMR** (500 MHz, $CDCl_3$) δ = 7.68 (1H, s), 7.25 (1H, t, J = 8.0 Hz), 7.14 (1H, d, J = 8.0 Hz), 7.06 (1H, s), 7.05 (1H, d) 5.07 (2H, s), 4.18 (2H, br s), 2.33 (3H, s); **$^{13}$C NMR** (125 MHz, $CDCl_3$) δ = 163.7, 142.2, 138.8, 134.7, 129.3, 128.9, 128.7, 125.0, 53.3, 21.3; **HRMS** (ESI-TOF) *m/z* $[M+H]^+$ calcd for $C_{10}H_{10}N_4O_2$ 188.1803, found 188.1805. **LCMS** RT= 0.48 min, *m/z* =188.83 $[M+H]^+$

**1-benzyl-1H-1,2,4-triazol-3-amine**

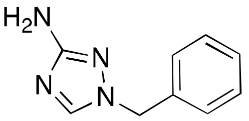


Prepared using Method B to give the title compound (300 mg, 1.72 mmol, 90 %) as an off-white solid. **Rf** 0.14 (1:1 petrol-EtOAc) **$^1$H NMR** (400 MHz, $CDCl_3$) $\delta$= 7.70 (1H, s), 7.40-7.34 (3H, m), 7.32-7.26 (2H, m), 5.13 (2H, s) 3.77 (2H, br s); **$^{13}$C NMR** (100 MHz, $CDCl_3$) $\delta$= 142.3, 134.8, 129.0, 128.5, 128.0, 53.3; **HRMS** ($ESI^+$) *m/z*: $[M + H]^+$ Calcd for $C_9H_{11}N_4$ 175.0983; Found 175.0974. **HPLC** RT = 1.09 min, 95.5 % relative area

**1-(4-chlorobenzyl)-1H-1,2,4-triazol-3-amine**

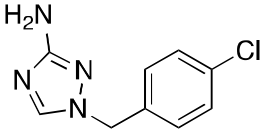


To a slurry of iron powder (655 mg, 11.73 mmol, 7.0 eq) in $MeOH/H_2O/AcOH$ (3 mL: 3 mL: 0.3 mL) was added 1-[(4-chlorophenyl)methyl]-3-nitro-1*H*-1,2,4-triazole (400 mg, 1.68 mmol, 1.0 eq). The reaction was refluxed at 80 °C for 1 h. The reaction was quenched with 2 M NaOH (3.0 mL) and filtered through celite, washing with MeOH. The filtrate was concentrated *in vacuo* and redissolved in EtOAc (50 mL). The organic solution was washed with water (50 mL) and extracted in EtOAc (3 x 50 mL), dried ($MgSO_4$) and concentrated to afford the title compound (85 mg, 0.41 mmol, 97 %) as a colourless solid. **Rf** 0.50 (5% MeOH in DCM); **$^1$H NMR** (400 MHz, $CDCl_3$) $\delta$= 7.81 (1H, s), 7.34 (1H, d, J= 8.0 Hz), 7.22 (1H, d, J= 8.0 Hz), 5.11 (2H, s), 3.55 (2H, br s); **$^{13}$C NMR** (400 MHz, $CDCl_3$) $\delta$= 163.7, 142.6, 134.2, 133.6, 129.2, 128.9, 52.1; **HRMS** ($ESI^+$) *m/z*: $[M + H]^+$ Calcd for $C_9H_{10}ClN_4$ 209.0594; Found 209.0583. **HPLC** RT = 1.52 min, 100 % relative area.

**8-Chloro-2-(4-pyridinyl)-4-quinolinecarboxylic acid**

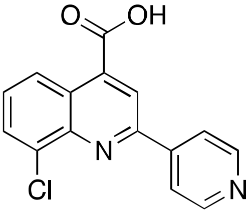


Prepared using Method C. Recrystallisation with methanol gave the title compound as an orange solid (400 mg, 1.41 mmol, 64%) **Rf** 0.43 (5 % MeOH in DCM);**$^1$H NMR** (400 MHz, DMSO-$d_6$) δ = 8.82 (2H, d, J = 7.5 Hz) 8.63 (2H, d, J = 7.0 Hz), 8.33 (2H, d, J = 7.0 Hz), 8.11 (1H, d, J = 7.5 Hz), 7.73 (1H, t, J = 8.0 Hz) **$^{13}$C NMR** (100 MHz, DMSO-$d_6$) δ = 167.6, 154.5, 151.7, 144.9, 133.9, 131.2, 129.2, 126.2, 125.4, 121.9, 120.3 **HRMS** (ESI-TOF) *m/z*: $[M + K]^+$ Calcd for $C_{15}H_9ClN_2O_2K$ 322.9984; Found 323.0012.; **HPLC** RT =1.79 min, (100% relative area).

**2-pyridyl-8-fluoroquinoline-4-carboxylic acid**

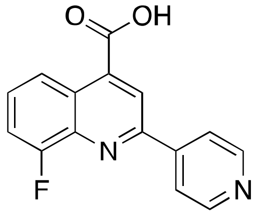


Prepared using Method C. Trituration with hot acetone gave <u>the title compound</u> as a dark orange solid (261mg, 0.973 mmol, 40%) which was used without further purification. **Rf** 0.00 (100% EtOAc); **$^1$H NMR** (500 MHz, DMSO-$d_6$) δ = 8.90 (2H, d, J = 7.5 Hz) 8.71 (1H, s), 8.52 (1H, d, J = 7.0 Hz), 8.30 (2H, d, J = 7.5 Hz), 7.73 (2H, d, J = 8.0 Hz) **$^{13}$C NMR** (125 MHz, DMSO-$d_6$) δ = 167.6, 160.0, 157.0, 154.3, 151.1, 150.8, 144.9, 139.1, 129.2, 126.2, 122.3, 122.0, 121.8, 120.6, 115.2 **HRMS** (ESI-TOF) *m/z* [M-H]$^-$ calcd for $C_{15}H_{10}FN_2O_2$ 269.072082, found 269.072132; **LCMS** RT = 0.43 min, m/z = 269.10 [M+H]$^+$.

**6-Methyl-2-(4-pyridinyl)-4-quinolinecarboxylic acid**

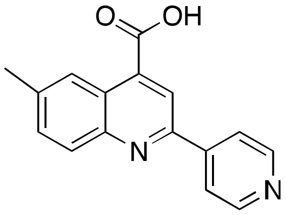


Prepared using Method C. Recrystallisation with methanol gave <u>the title compound</u> (340 mg, 1.19 mmol, 54 %) as a bright orange solid **Rf** 0.08 (5 % MeOH in DCM) **$^1$H NMR** (400 MHz, DMSO-$d_6$) *δ*= 8.79 (2H, d, J= 4.0 Hz), 8.52 (1H, s), 8.45 (1H, d, J= 1.0 Hz), 8.25 (2H, dd, $J_1$= 4.0 Hz, $J_2$ = 1.0 Hz), 8.13 (1H, d, J= 8.0 Hz), 7.76 (1H, dd, J= 8.6 Hz, J= 2.0 Hz), 2.58 (1H, s) **$^{13}$C NMR** (100 MHz, DMSO-$d_6$) δ = 167.8, 153.0, 150.9, 147.5, 145.4, 139.1, 133.1, 130.2, 124.6, 121.6, 119.5, 22.1*;* **HRMS (ESI$^+$)** *m/z*: [M + H]$^+$ Calcd for $C_{16}H_{13}N_2O_2$ 265.0977; Found 265.0971. **HPLC** RT = 1.70 min, 100 % relative area.

**2-Bromo-N-{1-[(3-methylphenyl)methyl]-1H-1,2,4-triazol-3-yl}pyridine-4-carboxamide**

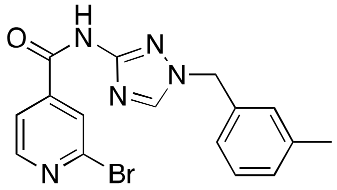


Prepared using Method D. Purification using column chromatography (4% MeOH in DCM) gave <u>the title compound</u> as a colourless solid (219 mg, 0.58 mmol, 59 %). **$^1$H NMR** (500 MHz, $CDCl_3$) δ = 10.90 (s, 1H), 8.45 (d, J = 4.97 Hz, 1H), 7.92 (s, 1H), 7.72 (1H, d, J = 4.97 Hz), 7.63 (1H, s), 7.27 (1H, t, J = 7.36 Hz), 7.18 (1H d, J = 7.36 Hz), 7.11 (1H, s), 7.09 (1H, d, J= 7.36 Hz), 5.28 (2H, s), 2.34 (3H, s); **$^{13}$C NMR** (125 MHz, $CDCl_3$) δ = 162.0, 156.0, 151.0, 144.0, 142.6, 141.9, 139.2, 133.2, 129.9, 129.4, 129.2, 126.0, 125.7, 121.2, 54.5, 21.4 **HRMS** (ESI-TOF) *m/z* [M+H]$^+$ calcd for $C_{16}H_{15}BrN_5O$ 372.0454, found 372.0455; **LCMS** RT = 0.5 min, *m/z* = 371.92 [M+H]$^+$

**2-(4-methylphenyl)-N-{1-[(3-methylphenyl)methyl]-1H-1,2,4-triazol-3-yl} quinoline-4-carboxamide.**

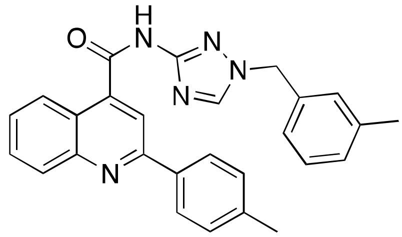


Prepared using Method D. Purification using column chromatography (0.5%-3% MeOH in DCM) gave <u>the title compound</u> as a colourless solid. (389 mg, 0.89 mmol, 45 %). **$^1$H NMR** (500 MHz, DMSO-$d_6$) δ = 11.22 (1H, s) 8.59 (1H, s), 8.19 (1H, s), 8.18 (2H, d, J = 8.0 Hz), 8.11 (1H, d, J = 6.5 Hz), 8.06 (1H, d, J = 8.5 Hz), 7.76 (1H, t, J = 8.0 Hz), 7.58 (1H, t, J = 8.0 Hz), 7.32 (2H, d, 8.0 Hz), 7.22 (1H, s), 7.11 (1H, d, J = 7.5 Hz) 7.10 (1H, t, J = 7.5 Hz), 7.09 (1H, d, J = 7.5 Hz), 5.31 (2H, s), 2.34 (3H, s), 2.25 (3H, s) **$^{13}$C NMR** (125 MHz, DMSO-$d_6$) δ = 165.3, 156.4, 156.2, 148.4, 144.5, 142.2, 140.1, 138.3, 136.5, 135.9, 130.7, 130.0, 129.3, 129.1, 129.1, 129.0, 127.7, 127.6, 125.6, 125.5, 123.6, 117.3, 52.9, 21.5, 21.4 **HRMS** (ESI-TOF) *m/z* [M+H]$^+$ calcd for $C_{27}H_{24}N_5O$ 433.1975, found 434.1979; **LCMS** RT = 0.7 min, *m/z* = 434.12 [M+H]$^+$.

**2-phenyl)-N-{1-[(3-methylphenyl)methyl]-1H-1,2,4-triazol-3-yl} quinoline-4-carboxamide**

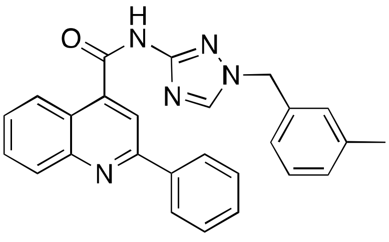

Prepared using Method D. Purification using column chromatography (4% MeOH in DCM) gave the title compound as a colourless solid (10mg, 0.0238 mmol, 7 %) **Rf** 0.24 (2 % MeOH in DCM) **$^{1}$H NMR** (400 MHz, $CDCl_3$) δ = 11.22 (1H, s) 8.32 (1H, d, J= 8.0 Hz), 8.19-8.05 (4H, m), 7.75 (1H, d, J = 8.0 Hz), 7.58-7.50 (4H, m), 7.24-7.18 (2H, m), 6.96-6.92 (2H, m), 6.75 (1H, br s), 5.08 (s, 2H), 2.37 (s, 3H) **$^{13}$C NMR** (100 MHz, $CDCl_3$) δ =156.5, 156.1, 148.7, 142.0, 141.3, 139.0, 138.5, 133.2, 130.3, 130.1, 129.7, 129.2, 127.5, 127.4, 125.5, 125.3, 123.2, 117.0, 54.2, 21.4 **HRMS** (ESI-TOF) *m/z* $[M+H]^+$ calcd for $C_{26}H_{22}N_5O$ 420.1819, found 420.1817; **HPLC** RT =2.88 min (100% relative area).

**8-chloro-N-(1-(3-tolyl)-1H-1,2,4-triazol-3-yl)-2-(pyridin-4-yl)quinoline-4-carboxamide**

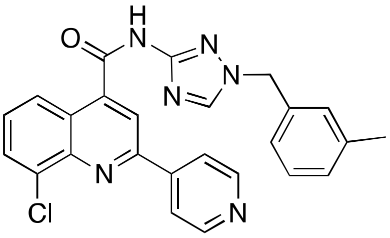

Prepared using Method D. Purification using column chromatography (1%-5% MeOH in DCM) gave the title compound as a peach solid. (25 mg, 0.055 mmol, 7 %). **Rf** 0.36 (5 % MeOH in DCM) **$^{1}$H NMR** (500 MHz, $CDCl_3$) δ = 10.71 (1H, s) 8.85 (2H, s), 8.23 (1H, s), 8.15 (1H, d, J = 8.1 Hz), 7.95 (1H, s), 7.54 (1H, s) 7.20-7.14 (4H, m) 6.96 (2H, br s), 5.15 (2H, s), 2.34 (3H, s) **$^{13}$C NMR** (125 MHz, $CDCl_3$) δ = 155.9, 153.9, 150.7, 145.0, 142.9, 141.6, 139.1, 134.9, 133.2, 131.0, 129.8, 129.6, 129.1, 128.3, 125.4, 124.3, 121.4, 117.0, 54.3, 21.3 **HRMS** (ESI-TOF) *m/z* $[M+H]^+$ calcd for $C_{25}H_{19}ClN_6O$ 455.1381, found 455.1377 **LCMS** RT = 0.62 min, *m/z* = 455.1 $[M+H]^+$.

**N-(1-benzyl-1H-1,2,4-triazol-3-yl)-2-chloroquinoline-4-carboxamide**

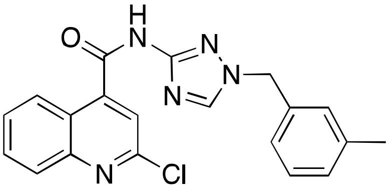

Prepared using Method D. Purification using column chromatography (1:1 Petrol/ EtOAc) gave the title compound as a colourless solid. (154 mg, 0.408 mmol, 41 %). **Rf** 0.33 (1:1 Petrol/ EtOAc) **$^{1}$H NMR** (400 MHz $CDCl_3$)δ = 11.22 (s, 1H), 8.22 (1H, d, J= 8.0 Hz), 7.95 (1H, d, J= 8.0 Hz), 7.70-7.65 (1H, m), 7.53-7.45 (2H, m), 7.24-7.10 (2H, m), 7.00-6.93 (3H, m), 5.14 (2H, s), 2.32 (3H, s) **$^{13}$C NMR** (100 MHz, $CDCl_3$) δ = 149.8, 148.4, 140.6, 139.3, 132.5, 131.4, 130.2, 129.5, 129.3, 128.9, 128.3, 125.9, 125.5, 123.2, 120.6, 54.9, 21.4; **HRMS** (ESI-TOF) *m/z* $[M+Na]^+$ calcd for $C_{20}H_{16}ClN_5NaO$ 400.093559, found; 400.094192 **LCMS** RT = 0.55 min, *m/z* = 377.90 $[M+H]^+$.

**8-fluoro-N-(1-(3-tolyl)-1H-1,2,4-triazol-3-yl)-2-(3-bromophenyl)quinoline-4-carboxamide**

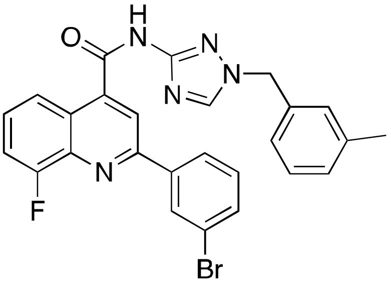

Prepared using Method D. Purification using column chromatography (100% EtOAc) gave the title compound as an off-white solid. (15 mg, 0.029 mmol, 7 %). **$^{1}$H NMR** (500 MHz, DMSO-$d_6$) δ = 11.3 (1H, s) 8.70-8.35 (4H, m), 8.05 (1H, d, J= 8.0 Hz), 7.75-7.60 (3H, m), 7.50 (1H, d, J = 8.0 Hz), 7.27-7.23 (1H, m), 7.20-7.10 (3H, m), 5.40 (2H, s), 2.30 (3H, s); **$^{13}$C NMR** (100 MHz, DMSO-$d_6$) δ = 144.4, 133.4, 131.6, 130.4, 130.3, 129.4, 129.1, 127.0, 125.5, 123.0, 115.1, 21.4 **$^{19}$F NMR** (400 MHz, DMSO-$d_6$) δ = -123.9 **HRMS** (ESI-TOF) *m/z* $[M+H]^+$ calcd for $C_{26}H_{20}BrFN_5O$ 516.082977, found 516.083847; **LCMS** RT = 0.7 min, *m/z* = 517.03 $[M+H]^+$.

**N-2-Benzothiazolyl-2-(4-methylphenyl)-4-quinolinecarboxamide**

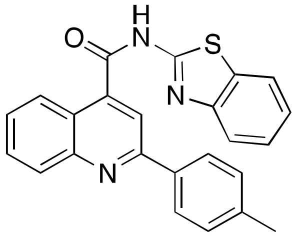

Prepared using Method D. Purification using column chromatography (3:1 Petrol/EtOAc) gave the title compound as an off-white solid. (58 mg, 1.47 mmol, 85 %) **Rf** 0.70 (3:1 Petrol/EtOAc) **$^1$H NMR** (400 MHz $CDCl_3$)δ = 8.60 (2H, br s) 8.10 (1H, s), 7.89 (2H, d, J= 8.0 Hz), 7.80-7.65 (2H, m), 7.58 (1H, t, J = 8.0 Hz), 7.48- 7.28 (3H, m), 7.17 (2H, d, J= 8.0 Hz) 2.23 (s, 3H); **$^{13}$C NMR** (400 MHz $CDCl_3$) δ= 165.0, 129.7, 128.2, 126.9, 124.5, 121.7, 21.3 **HRMS** (ESI-TOF) *m/z* $[M+H]^+$ calcd for $C_{24}H_{18}N_3OS$ 396.116510, found 396.117665; **LCMS** RT = 0.76 min, *m/z* = 396.34 $[M+H]^+$.

***N*-1H-benzotriazo-1-yl-2-(4-tolyl)-4-quinolinecarboxamide**

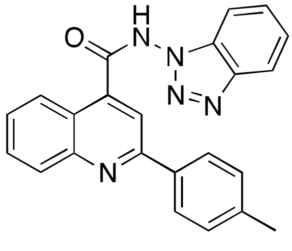

Prepared using Method D. Purification using column chromatography (3:1 Petrol/EtOAc) gave the title compound as an off-white solid. (8 mg, 0.021 mmol, 11%) **Rf** 0.22 (3:1 Petrol/EtOAc) **$^1$H NMR** (400 MHz $CDCl_3$)δ = 8.83 (1H, d, J= 8.0 Hz) 8.63 (1H, s), 8.46 (1H, d, J= 8.0 Hz), 8.18-8.08 (2H, m), 8.05 (1H, d, J = 8.0 Hz), 7.80 (1H, t, J= 8.0 Hz) 7.70- 7.55 (2H, m) 7.52-7.40 (3H, m), 7.39-7.30 (2H, m) 7.07 (2H, d, J=8.0 Hz) 5.68 (2H, br s) 2.25 (s, 3H) **LCMS** RT = 0.65 min, *m/z* = 380.31 $[M+H]^+$.

***N*-(1-benzyl-1H-1,2,4-triazol-3-yl)-2-(p-tolyl)quinoline-4-carboxamide**

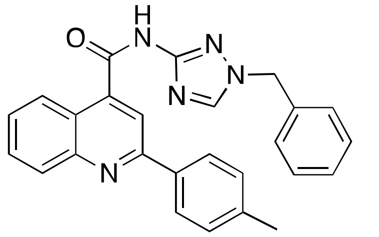

Prepared using Method D. Purification using column chromatography (2 % MeOH in DCM) gave the title compound (86 mg, 0.21 mmol, 36 %) as a light brown solid **Rf** 0.24 (5 % MeOH in DCM) **$^1$H NMR** (500 MHz, MeOD-$d_4$) *δ*= 8.56 (1H, s), 8.19 (2H, d, J= 8.8 Hz), 8.11 (1H, d, J= 8.8 Hz), 7.77 (2H, d, J= 4.0 Hz), 7.59 (1H, t, J= 7.5 Hz), 7.31 (7H, m), 5.38 (2H, s), 2.38 (3H, s) **$^{13}$C NMR (**125 MHz, MeOD-$d_4$) *δ*= 156.3, 144.1,140.0, 136.3, 135.9, 130.3, 129.7, 128.85, 128.81, 128.2, 128.1, 127.9, 125.3, 123.4, 117.2, 52.9, 20.7; **HRMS (ESI$^+$)** *m/z*: $[M + H]^+$ Calcd for $C_{26}H_{22}N_5O$ 420.1824; Found 420.1744. **HPLC** RT = 2.71 min, 100 % relative area.

***N*-(1-benzyl-1H-1,2,4-triazol-3-yl)-8-chloro-2-methylquinoline-4-carboxamide**

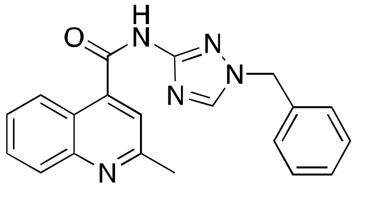

Prepared using Method D. Purification using column chromatography (3 % - 10 % MeOH in DCM) gave the title compound (101 mg, 0.29 mmol, 51 %) as a light brown solid **Rf** 0.10 (5 % MeOH in DCM); **$^1$H NMR (**600 MHz, MeOD-$d_4$) *δ*= 8.42 (1H, s), 8.19 (1H, d, J= 8.4 Hz), 8.00 (1H, d, J= 8.4 Hz), 7.76 (1H, t, J = 7.5 Hz), 7.58 (2H, d, J= 8.8 Hz), 7.36 (3H, m), 7.32 (2H, m), 5.39 (2H, s), 2.75 (3H, s) **$^{13}$C NMR** (150 MHz, MeOD-$d_4$) δ= 180.6, 143.6, 135.4, 130.2, 128.6, 128.5, 128.1, 128.0, 127.9, 127.7, 127.5, 126.9, 124.9, 122.8, 120.1, 53.2, 23.4; **HRMS (ESI$^+$)** *m/z*: $[M + H]^+$ Calcd for $C_{20}H_{18}N_5O$ 344.1511; Found 344.1501. **HPLC** RT = 1.54 min, 100 % relative area.

**8-chloro-N-(1-(4-chlorobenzyl)-1H-1,2,4-triazol-3-yl)-2-(pyridin-4-yl)quinoline-4-carboxamide**

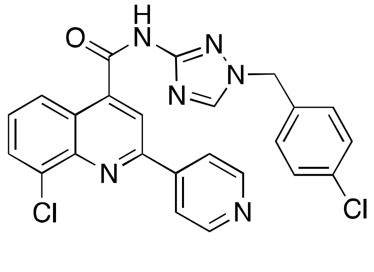

Prepared using Method D. Purification using column chromatography (5 % MeOH in DCM) gave the title compound (15 mg, 0.03 mmol, 8 %) as a red solid. **Rf** 0.10 (5 % MeOH in DCM); **$^1$H NMR** (400 MHz, $CDCl_3$) δ 8.68 (1H, dd, $J_1$ =4.8 Hz, $J_2$ =1.4 Hz), 8.26 (1H, s), 8.19 (3H, d, J = 4.0 Hz), 7.94 (1H, d), 7.88 (1H, d, J= 8.0 Hz), 7.51 (1H, t, J= 8.0 Hz), 7.31 (1H, s), 7.25 (3H, s), 5.29 (2H, s) **$^{13}$C NMR** (100 MHz, $CDCl_3$) δ **=** 160.0, 153.8, 150.1,

146.8, 138.6, 134.7, 133.5, 133.1, 132.2, 125.9, 57.3 **HRMS (ESI$^+$)** *m/z*: [M + H]$^+$ Calcd for $C_{24}H_{17}Cl_2N_6O$ 475.0841; Found 475.0834. **HPLC** RT = 2.34 min, 100 % relative area.

**6-methyl-N-(1-(3-methylbenzyl)-1H-1,2,4-triazol-3-yl)-2-(pyridin-4-yl)quinoline-4-carboxamide**

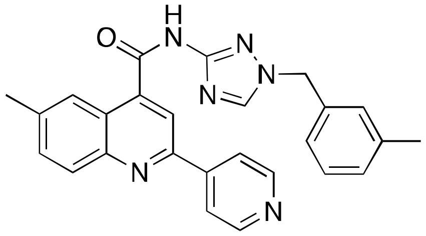

Prepared using Method D. Purification using column chromatography (2-20% MeOH in DCM) gave the title compound (86 mg, 0.21 mmol, 36%) as a light brown solid **Rf** 0.42 (5% MeOH in DCM) **$^1$H NMR** (500 MHz, MeOD-$d_4$) $\delta$= 8.62 (2H, d, J = 5.5 Hz), 8.37 (1H, s), 8.15-8.21 (3H, m), 8.03 (1H, d, J= 8.8 Hz), 7.99 (1H, s), 7.62 (1H, d, J = 8.8 Hz) 7.15 (1H, t, J= 7.0 Hz), 7.13 (1H, s), 7.08 (2H, t, J= 7.0 Hz), 5.28 (2H, s), 2.47 (3H, s), 2.25 (3H, s) **$^{13}$C NMR** (125 MHz, MeOD-$d_4$) $\delta$= 168.1, 157.9, 154.4, 151.4, 149.1, 148.7, 145.5, 143.5, 140.9, 140.5, 137.2, 134.7, 133.6, 131.4, 130.8, 130.5, 128.9, 126.9, 126.0, 125.6, 123.7, 118.7, 55.1, 22.6 **HRMS (ESI$^+$)** *m/z:* [M + H]$^+$ Calcd for $C_{26}H_{22}N_6O$ 435.1933; Found 435.1924. **HPLC** RT = 2.23 min, 100 % relative area.

***N*-{1-[(3-Methylphenyl)methyl]-1H-1,2,4-triazol-3-yl}-1H-indole-3-carboxamide**

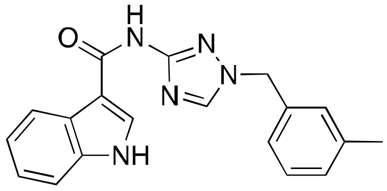

3-Indole-carboxylic acid (100 mg, 0.621 mmol, 1.0 eq.) was cooled to 0 °C, and $SOCl_2$ (0.5 mL) added dropwise. The reaction was stirred and allowed to warm to room temperature, and then refluxed for 1 hr, after which no solids were present. The reaction was concentrated to yield a pink waxy solid, to which was added dropwise a solution of 1-[(3-methylphenyl)methyl]-1H-1,2,4-triazol-3-amine (128 mg, 0.681 mmol, 1.1 eq.) and $NEt_3$ (0.24 mL, 1.86 mmol, 3.0 eq.) in DCM (5 mL). The reaction was allowed to stir at room temperature for 18 h, after which a white precipitate had formed. The solids were filtered, triturated in diethyl ether, and dried under vacuum to yield the title compound as a colourless solid (152 mg, 0.459 mmol, 74%). **$^1$H NMR** (500 MHz, DMSO) δ = 11.70 (1H, s) 10.19 (1H, s), 8.56 (1H, s), 8.30 (1H, d, J = 3.0 Hz), 8.16, (1H, d, J = 7.5 Hz), 7.45 (1H, d, J = 7.5 Hz), 7.27 (1H, t, J = 7.5 Hz), 7.18 (1H, dt, $J_1$ = 8.0 Hz, $J_2$ = 1.5 Hz) 7.15 (1H, s), 7.14 (1H, dt, $J_1$ = 8.0 Hz, $J_2$ = 1.5 Hz) 7.14 (1H, s), 7.13 (1H, d, J = 7.5 Hz) 5.32 (2H, s) 2.31 (3H, s) **$^{13}$C NMR** (125 MHz, DMSO-$d_6$) δ = 163.1, 157.2, 144.1, 138.3, 136.7, 136.6, 129.6, 129.0, 128.9, 127.0, 125.5, 122.6, 121.5, 121.2, 116.0, 112.4, 110.1, 52.7, 21.4; **HRMS** (ESI-TOF) *m/z* [M+H]$^+$ calcd for $C_{19}H_{18}N_5O$ 332.1506, found 332.1508; **LCMS** RT = 0.6 min, *m/z* = 332.04 [M+H].

**1-(4-Methylphenyl)-N-{1-[(3-methylphenyl)methyl]-1H-1,2,4-triazol-3-yl}-1H-indole-3-carboxamide**

N-{1-[(3-methylphenyl)methyl]-1H-1,2,4-triazol-3-yl}-1H-indole-3-carboxamide (75 mg, 0.23 mmol, 1.0 eq.), 4-bromo-toluene (37 mg, 0.22 mmol, 1.2 eq.), copper iodide (8 mg, 0.04 mmol, 0.2 eq.) and $K_3PO_4$ (75 mg, 0.36 mmol, 2.1 eq.) were placed under $N_2$ atmosphere. Anhydrous DMF (2.5 mL) as added and the reaction degassed for 10 minutes. To this was added a solution of N,N′-dimethylethylenediamine (8 µL, 0.07 mmol, 0.4 eq.) in anhydrous DMF (0.5 mL), upon which the reaction instantaneously went from colourless to black. The reaction was heated to 110 °C for 18 h. After this, the reaction was judged to have not gone to completion, so was cooled to room temperature, and further 4-bromo-toluene (40 mg, 0.23 mmol, 1.3 eq.), copper iodide (20 mg, 0.11 mmol, 0.6 eq.) $K_3PO_4$ (35 mg, 0.17 mmol, 0.9 eq.) and N,N′-dimethylethylenediamine (20 µL, 0.18 mmol, 1.0 eq.) was added under $N_2$. The reaction was heated to 110 °C for 18 h. The reaction was cooled to room temperature, filtered through silica plug (eluted with 10% MeOH in DCM), and filtrate concentrated to give a pale green solid. This was purified by column chromatography (3% MeOH in DCM), and relevant fractions concentrated. The residue was dissolved in DCM (20 mL), washed with 10% $Na_2SO_4$ (2 x 10 mL), dried ($Na_2SO_4$) and concentrated to yield the title compound as a pale yellow solid (16 mg, 38.0 µmol, 17 %). **$^1$H NMR** (500 MHz, DMSO-$d_6$) 10.39 (1H, s) 8.56 (1H, s) 8.58 (1H, s), 8.31-8.27 (1H, m) 7.55-7.68 (1H, m), 7.60-7.54 (1H, m), 7.53-7.50 (1H, m) 7.45 (2H, d, J= 8.0 Hz) 7.30-7.26 (3H, m), 7.12 (1H, t, J = 7.5 Hz), 7.16 (1H, s), 7.14 (1H, t, J = 8.0 Hz), 5.33 (2H, s), 2.43 (3H, s), 2.31 (3H, s) **$^{13}$C NMR** (125 MHz, DMSO-$d_6$) δ = 162.4, 157.0, 144.1, 138.3, 138.2, 137.5, 136.7, 136.2, 136.2, 132.3, 130.9, 129.0, 129.0, 127.9, 125.5, 124.6, 123.8, 122.4, 122.1, 111.3, 111.2, 52.8, 21.4, 21.1 **HRMS** (ESI-TOF) *m/z* $[M+H]^+$ calcd for $C_{26}H_{24}N_5O$ 422.1979, found 422.1980. **LCMS** RT = 0.8 min, *m/z* = 422.09 $[M+H]^+$

**2-(4-Methylphenyl)-N-{1-[(3-methylphenyl)methyl]-1H-1,2,4-triazol-3-yl}pyridine-4-carboxamide**

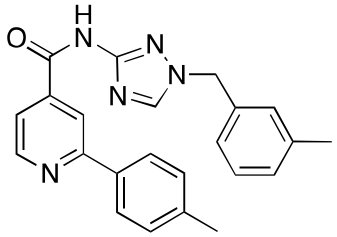


Prepared using Method E. Purification using column chromatography (5 % MeOH in DCM) gave the title compound as a colourless solid (24 mg, 62.6 µmol, 51 %) **$^1$H NMR** (500 MHz, $CDCl_3$) δ = 9.50 (1H, s), 8.83 (1H, d, J = 5.09 Hz), 8.18 (1H,s), 7.95 (1H, d, J = 7.5 Hz), 7.73 (1H, s), 7.64 (1H, d, J = 5.0 Hz) 7.33 (2H, d) 7.28 (1H t, J = 8.0 Hz) 7.19 (1H, d, J = 8.0 Hz), 7.10 (1H, s), 7.09 (1H, d, J = 8.0 Hz), 5.26 (2H, s), 2.45 (3H, s), 2.37 (3H, s) **$^{13}$C NMR** (125 MHz, $CDCl_3$) δ 163.3, 158.6, 156.0, 150.5, 142.1, 141.8, 139.9, 139.0, 135.5, 133.5, 129.7, 129.7, 129.2, 129.1, 126.9, 125.5, 119.1, 117.8, 54.3, 21.3, 21.3 **HRMS** (ESI-TOF) *m/z* $[M+H]^+$ calcd for $C_{23}H_{22}N_5O$ 384.1817, found 384.1818; **LCMS** RT = 0.7 min, *m/z* = 384.35 $[M+H]^+$

**Small molecule crystal structure of 1-[(3-Methylphenyl)methyl]-3-nitro-1H-1,2,4-triazole**

Measurements were carried out at 125K on an Agilent SuperNova diffractometer equipped with an Atlas CCD detector and connected to an Oxford Cryostream low temperature device using mirror monochromated Cu $K_\alpha$ radiation (λ = 1.54184 Å) from a Microfocus X-ray source. The structure was solved by intrinsic phasing using SHELXT[11] and refined by a full matrix least squares technique based on $F^2$ using SHELXL2014.[12]

The compound crystallised as colourless blocks. The compound crystallised in a monoclinic cell and was solved in the $P2_1/c$ space group, with two molecules in the asymmetric unit. All non-hydrogen atoms were located in the Fourier Map and refined anisotropically. All hydrogen atoms were placed in calculated positions and refined isotropically using a "riding model". Pictures are presented with non-hydrogen atoms displayed as displacement ellipsoids, which are set at the 50% probability level.

Molecular Formula: $C_{10}H_{10}N_4O_2$

Molecular structure:

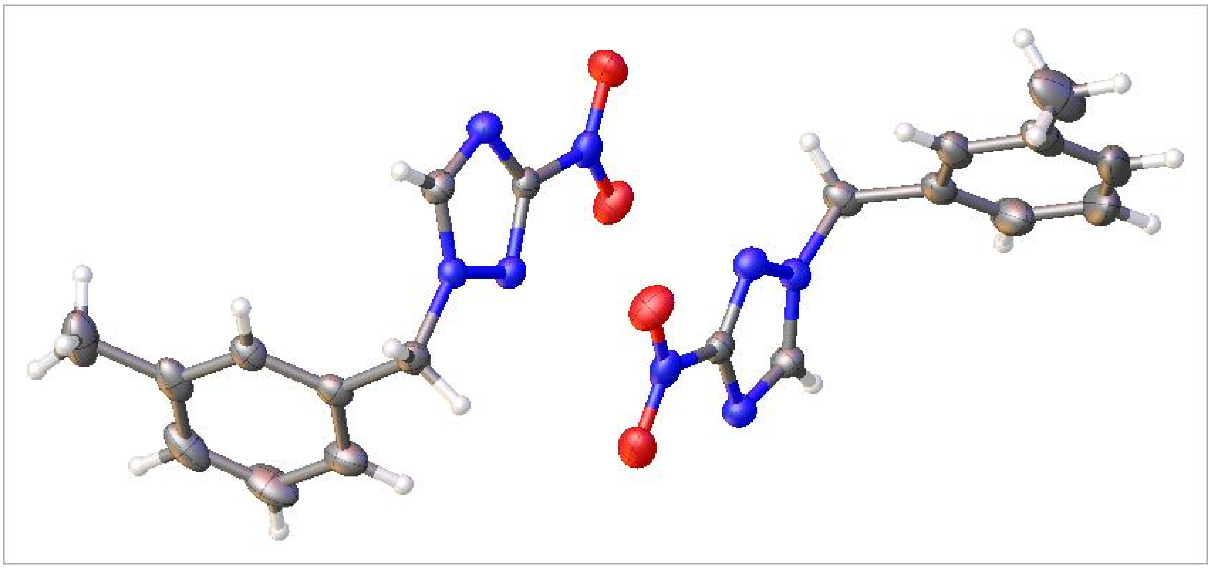

**Figure S4-** Small molecule crystal structure of 1-[(3-Methylphenyl)methyl]-3-nitro-1H-1,2,4-triazole

**Table S5**: **Crystal data and structure refinement for 1-[(3-Methylphenyl)methyl]-3-nitro-1H-1,2,4-triazole**

| **Identification code** | **MJM424** |
|---|---|
| **Empirical formula** | $C_{10}H_{10}N_4O_2$ |
| **Formula weight** | 218.22 |
| **Temperature/K** | 125.00(10) |
| **Crystal system** | monoclinic |
| **Space group** | $P2_1/c$ |
| **a/Å** | 13.5667(2) |
| **b/Å** | 7.95929(14) |
| **c/Å** | 19.6589(3) |
| **α/°** | 90 |
| **β/°** | 93.4835(14) |
| **γ/°** | 90 |
| **Volume/Å$^3$** | 2118.88(6) |
| **Z** | 8 |
| **$\rho_{calc}$g/cm$^3$** | 1.368 |
| **μ/mm$^{-1}$** | 0.833 |
| **F(000)** | 912.0 |
| **Crystal size/mm$^3$** | 0.19 × 0.17 × 0.08 |
| **Radiation** | CuKα (λ = 1.54184) |
| **2Θ range for data collection/°** | 10.81 to 147.866 |
| **Index ranges** | $-12 \leq h \leq 16$, $-7 \leq k \leq 9$, $-24 \leq l \leq 22$ |
| **Reflections collected** | 8178 |
| **Independent reflections** | 4141 [$R_{int}$ = 0.0244, $R_{sigma}$ = 0.0320] |
| **Data/restraints/parameters** | 4141/0/291 |
| **Goodness-of-fit on F$^2$** | 1.040 |
| **Final R indexes [I>=2σ (I)]** | $R_1$ = 0.0459, $wR_2$ = 0.1195 |
| **Final R indexes [all data]** | $R_1$ = 0.0537, $wR_2$ = 0.1273 |

| Largest diff. peak/hole / e Å$^{-3}$ | 0.58/-0.26 |
|---|---|

**Table S6**: **Fractional Atomic Coordinates (×104) and Equivalent Isotropic Displacement Parameters (Å2×103) for 1-[(3-Methylphenyl)methyl]-3-nitro-1H-1,2,4-triazole. Ueq is defined as 1/3 of of the trace of the orthogonalised UIJ tensor**

| Atom | *x* | *y* | *z* | U(eq) |
|---|---|---|---|---|
| **O1** | 2711.6(9) | 3881.2(17) | 2117.1(7) | 40.1(3) |
| **O2** | 1879.6(9) | 4977.7(17) | 2916.8(7) | 41.3(3) |
| **O3** | 3138.9(8) | 10130.0(16) | 2460.0(7) | 36.9(3) |
| **O4** | 2095.6(10) | 11079.7(17) | 3158.8(7) | 39.7(3) |
| **N1** | 4328.1(10) | 7646.8(17) | 3311.8(7) | 24.4(3) |
| **N2** | 3416.0(10) | 6965.2(17) | 3304.3(7) | 26.7(3) |
| **N3** | 4314.1(10) | 5786.2(18) | 2488.3(7) | 27.8(3) |
| **N4** | 2621.7(10) | 4835.9(18) | 2598.1(7) | 29.9(3) |
| **N5** | 858.1(9) | 7319.8(16) | 1761.7(6) | 22.3(3) |
| **N6** | 1766.0(9) | 7981.6(17) | 1911.2(7) | 24.1(3) |
| **N7** | 608.2(9) | 9283.7(18) | 2515.9(7) | 26.9(3) |
| **N8** | 2320.1(10) | 10190.2(17) | 2683.1(7) | 27.6(3) |
| **C1** | 4408.6(15) | 7561(2) | 4926.2(9) | 35.5(4) |
| **C2** | 4754.0(19) | 6899(2) | 5555.2(9) | 46.7(5) |
| **C3** | 5754(2) | 6896(3) | 5706.7(12) | 63.4(8) |
| **C4** | 6404(2) | 7518(4) | 5260.0(14) | 71.5(9) |
| **C5** | 6052.7(16) | 8186(3) | 4636.0(11) | 52.5(6) |
| **C6** | 5046.2(14) | 8209(2) | 4470.7(8) | 32.5(4) |
| **C7** | 4640.5(14) | 8948(2) | 3805.4(8) | 32.2(4) |
| **C8** | 4846.2(11) | 6940(2) | 2827.8(8) | 26.6(3) |
| **C9** | 3466.6(11) | 5883(2) | 2801.5(8) | 23.6(3) |
| **C10** | 4056(2) | 6236(3) | 6035.2(11) | 69.1(8) |
| **C11** | -186.6(13) | 7329(2) | 252.1(8) | 29.3(4) |
| **C12** | -251.9(15) | 7965(2) | -409.6(9) | 36.4(4) |
| **C13** | 586.1(17) | 7881(3) | -780.3(9) | 45.6(5) |
| **C14** | 1446.9(17) | 7188(3) | -508.1(10) | 50.0(6) |
| **C15** | 1501.9(14) | 6549(3) | 154.9(9) | 39.6(4) |
| **C16** | 674.2(12) | 6623(2) | 538.4(8) | 27.1(3) |
| **C17** | 706.4(13) | 5965(2) | 1258.1(8) | 28.2(4) |
| **C18** | 186.3(11) | 8104(2) | 2121.7(8) | 26.0(3) |
| **C19** | 1551.1(11) | 9131.4(19) | 2362.6(7) | 22.7(3) |
| **C20** | -1198.8(17) | 8704(3) | -713.7(11) | 51.9(6) |

**Table S7: Anisotropic Displacement Parameters (Å2×103) for 1-[(3-Methylphenyl)methyl]-3-nitro-1H-1,2,4-triazole. The Anisotropic displacement factor exponent takes the form: -2π2[h2a*2U11+2hka*b*U12+…]**

| Atom | $U_{11}$ | $U_{22}$ | $U_{33}$ | $U_{23}$ | $U_{13}$ | $U_{12}$ |
|---|---|---|---|---|---|---|
| **O1** | 42.6(7) | 31.3(7) | 45.0(7) | -6.1(6) | -9.8(6) | -0.7(6) |
| **O2** | 27.3(6) | 37.9(8) | 59.3(9) | 6.1(6) | 6.6(6) | -3.4(5) |

| | | | | | | |
|---|---|---|---|---|---|---|
| **O3** | 25.9(6) | 30.2(7) | 54.2(8) | 3.3(6) | -1.4(5) | -2.2(5) |
| **O4** | 45.5(7) | 34.7(7) | 37.8(7) | -11.6(6) | -6.7(6) | -0.4(6) |
| **N1** | 27.0(6) | 23.8(7) | 22.6(6) | 1.7(5) | 2.2(5) | -0.5(5) |
| **N2** | 27.9(7) | 25.2(7) | 27.2(7) | 3.1(5) | 4.3(5) | 0.7(5) |
| **N3** | 28.0(7) | 27.6(7) | 27.8(7) | -0.7(6) | 2.6(5) | 2.3(6) |
| **N4** | 28.4(7) | 23.1(7) | 37.3(8) | 4.9(6) | -4.5(6) | 1.7(6) |
| **N5** | 24.4(6) | 21.6(7) | 21.0(6) | 1.9(5) | 0.8(5) | -0.4(5) |
| **N6** | 23.0(6) | 22.5(7) | 26.7(6) | 1.5(5) | 0.9(5) | 0.6(5) |
| **N7** | 26.3(7) | 26.7(7) | 28.0(7) | -1.1(6) | 3.7(5) | 1.1(5) |
| **N8** | 28.0(7) | 21.7(7) | 32.3(7) | 3.8(6) | -5.5(5) | 0.7(5) |
| **C1** | 50.0(11) | 27.6(9) | 28.2(8) | -2.0(7) | -3.8(7) | 1.3(8) |
| **C2** | 87.3(16) | 24.4(9) | 27.6(9) | -4.8(7) | -3.9(9) | 7.2(10) |
| **C3** | 92.1(19) | 52.5(14) | 41.5(12) | -13.5(11) | -28.8(13) | 35.4(13) |
| **C4** | 57.5(14) | 93(2) | 60.3(15) | -32.6(15) | -26.1(12) | 37.1(15) |
| **C5** | 42.0(11) | 66.6(15) | 48.1(12) | -22.2(11) | -2.1(9) | 8.8(10) |
| **C6** | 42.0(9) | 27.4(9) | 27.6(8) | -7.7(7) | -2.6(7) | 1.7(7) |
| **C7** | 42.9(9) | 26.7(9) | 27.2(8) | -2.3(7) | 3.3(7) | -5.2(7) |
| **C8** | 23.6(7) | 27.9(8) | 28.5(8) | -0.1(7) | 3.5(6) | 1.7(6) |
| **C9** | 24.5(7) | 22.0(8) | 24.1(7) | 4.4(6) | -0.1(6) | 3.0(6) |
| **C10** | 124(2) | 47.9(14) | 35.5(11) | 6.3(10) | 9.2(13) | -7.2(15) |
| **C11** | 35.9(9) | 25.7(8) | 26.4(8) | -1.3(7) | 2.6(6) | -7.4(7) |
| **C12** | 52.3(11) | 27.5(9) | 28.3(8) | 0.4(7) | -6.2(8) | -14.0(8) |
| **C13** | 64.6(13) | 48.6(12) | 23.8(8) | -1.8(8) | 3.9(8) | -24.2(10) |
| **C14** | 52.6(12) | 65.4(15) | 34.1(10) | -12.3(10) | 21.0(9) | -17.9(11) |
| **C15** | 36.0(9) | 47.2(11) | 36.3(9) | -12.2(8) | 7.1(7) | -4.1(8) |
| **C16** | 34.1(8) | 22.7(8) | 24.7(8) | -4.8(6) | 2.7(6) | -6.1(6) |
| **C17** | 35.3(8) | 21.6(8) | 27.2(8) | -1.7(6) | -1.1(6) | -0.7(7) |
| **C18** | 23.1(7) | 26.5(8) | 28.6(8) | 0.9(6) | 2.6(6) | -0.5(6) |
| **C19** | 23.5(7) | 21.4(8) | 22.9(7) | 3.6(6) | -1.9(5) | 0.9(6) |
| **C20** | 68.0(14) | 41.4(12) | 43.5(11) | 12.0(9) | -18.7(10) | -10.3(10) |

**Table S8: Bond Lengths for 1-[(3-Methylphenyl)methyl]-3-nitro-1H-1,2,4-triazole.**

| Atom | Atom | Length/Å | | Atom | Atom | Length/Å |
|---|---|---|---|---|---|---|
| **O1** | N4 | 1.2250(19) | | N8 | C19 | 1.455(2) |
| **O2** | N4 | 1.2228(19) | | C1 | C2 | 1.398(3) |
| **O3** | N8 | 1.2198(18) | | C1 | C6 | 1.382(3) |
| **O4** | N8 | 1.2259(19) | | C2 | C3 | 1.371(4) |
| **N1** | N2 | 1.3503(19) | | C2 | C10 | 1.475(3) |
| **N1** | C7 | 1.464(2) | | C3 | C4 | 1.374(4) |
| **N1** | C8 | 1.341(2) | | C4 | C5 | 1.394(4) |
| **N2** | C9 | 1.316(2) | | C5 | C6 | 1.384(3) |
| **N3** | C8 | 1.323(2) | | C6 | C7 | 1.507(2) |
| **N3** | C9 | 1.339(2) | | C11 | C12 | 1.393(2) |
| **N4** | C9 | 1.453(2) | | C11 | C16 | 1.384(2) |
| **N5** | N6 | 1.3554(18) | | C12 | C13 | 1.389(3) |
| **N5** | C17 | 1.470(2) | | C12 | C20 | 1.504(3) |

| | | | | | | |
|---|---|---|---|---|---|---|
| **N5** | C18 | 1.341(2) | | C13 | C14 | 1.370(3) |
| **N6** | C19 | 1.320(2) | | C14 | C15 | 1.397(3) |
| **N7** | C18 | 1.325(2) | | C15 | C16 | 1.391(2) |
| **N7** | C19 | 1.337(2) | | C16 | C17 | 1.507(2) |

**Table S9: Bond Angles for 1-[(3-Methylphenyl)methyl]-3-nitro-1H-1,2,4-triazole**.

| Atom | Atom | Atom | Angle/° | | Atom | Atom | Atom | Angle/° |
|---|---|---|---|---|---|---|---|---|
| **N2** | N1 | C7 | 121.31(13) | | C1 | C6 | C5 | 119.35(19) |
| **C8** | N1 | N2 | 110.07(13) | | C1 | C6 | C7 | 119.86(16) |
| **C8** | N1 | C7 | 128.61(14) | | C5 | C6 | C7 | 120.78(18) |
| **C9** | N2 | N1 | 100.47(12) | | N1 | C7 | C6 | 112.01(14) |
| **C8** | N3 | C9 | 100.70(13) | | N3 | C8 | N1 | 110.84(14) |
| **O1** | N4 | C9 | 117.00(14) | | N2 | C9 | N3 | 117.92(14) |
| **O2** | N4 | O1 | 125.20(15) | | N2 | C9 | N4 | 120.37(14) |
| **O2** | N4 | C9 | 117.80(14) | | N3 | C9 | N4 | 121.71(14) |
| **N6** | N5 | C17 | 121.29(13) | | C16 | C11 | C12 | 122.09(17) |
| **C18** | N5 | N6 | 110.03(13) | | C11 | C12 | C20 | 121.18(18) |
| **C18** | N5 | C17 | 128.67(13) | | C13 | C12 | C11 | 117.69(18) |
| **C19** | N6 | N5 | 100.38(12) | | C13 | C12 | C20 | 121.13(18) |
| **C18** | N7 | C19 | 100.89(13) | | C14 | C13 | C12 | 121.32(18) |
| **O3** | N8 | O4 | 124.81(14) | | C13 | C14 | C15 | 120.45(18) |
| **O3** | N8 | C19 | 117.80(14) | | C16 | C15 | C14 | 119.41(19) |
| **O4** | N8 | C19 | 117.39(13) | | C11 | C16 | C15 | 119.04(16) |
| **C6** | C1 | C2 | 121.6(2) | | C11 | C16 | C17 | 119.90(15) |
| **C1** | C2 | C10 | 120.4(2) | | C15 | C16 | C17 | 121.05(16) |
| **C3** | C2 | C1 | 117.8(2) | | N5 | C17 | C16 | 111.88(13) |
| **C3** | C2 | C10 | 121.7(2) | | N7 | C18 | N5 | 110.79(14) |
| **C2** | C3 | C4 | 121.7(2) | | N6 | C19 | N7 | 117.90(14) |
| **C3** | C4 | C5 | 120.1(2) | | N6 | C19 | N8 | 120.73(14) |
| **C6** | C5 | C4 | 119.4(2) | | N7 | C19 | N8 | 121.37(14) |

**Table S10: Hydrogen Atom Coordinates (Å×104) and Isotropic Displacement Parameters (Å2×103) for 1-[(3-Methylphenyl)methyl]-3-nitro-1H-1,2,4-triazole**.

| Atom | *x* | *y* | *z* | U(eq) |
|---|---|---|---|---|
| **H1** | 3733 | 7565 | 4812 | 43 |
| **H3** | 5998 | 6461 | 6122 | 76 |
| **H4** | 7079 | 7493 | 5374 | 86 |
| **H5** | 6491 | 8612 | 4333 | 63 |
| **H7A** | 4081 | 9662 | 3889 | 39 |
| **H7B** | 5143 | 9642 | 3615 | 39 |
| **H8** | 5492 | 7222 | 2742 | 32 |
| **H10A** | 3550 | 7055 | 6100 | 104 |
| **H10B** | 4403 | 5998 | 6464 | 104 |
| **H10C** | 3759 | 5223 | 5853 | 104 |
| **H11** | -739 | 7380 | 509 | 35 |

| | | | | |
|---|---|---|---|---|
| **H13** | 563 | 8303 | -1222 | 55 |
| **H14** | 1998 | 7144 | -767 | 60 |
| **H15** | 2087 | 6078 | 338 | 48 |
| **H17A** | 91 | 5392 | 1333 | 34 |
| **H17B** | 1238 | 5155 | 1321 | 34 |
| **H18** | -484 | 7853 | 2098 | 31 |
| **H20A** | -1724 | 7905 | -681 | 78 |
| **H20B** | -1125 | 8972 | -1184 | 78 |
| **H20C** | -1352 | 9707 | -470 | 78 |

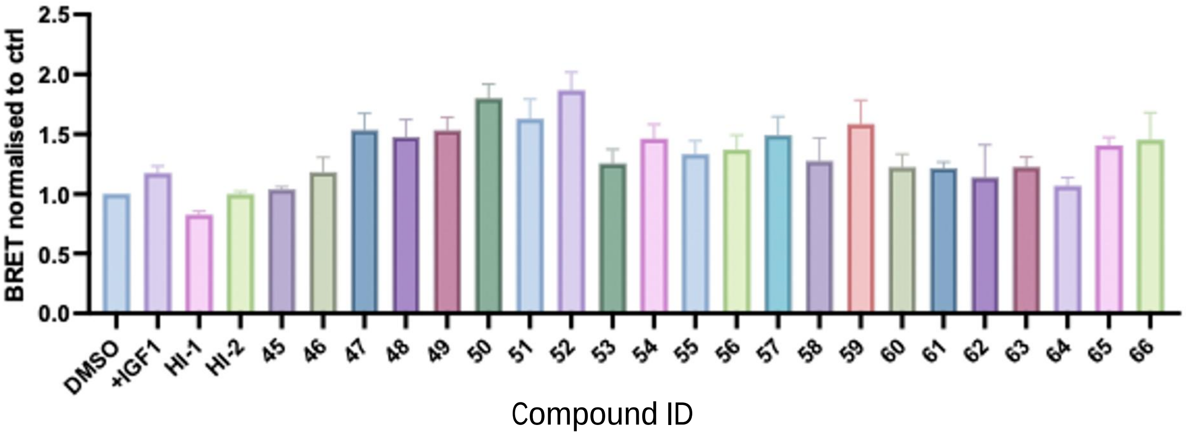


**Figure S5**: Compounds identified as analogues of HI-1 and screened using the BRET assay. Compound structures are displayed in Table S1. All compounds were screened at 100 µM final concentration (n=8,3).
